# HIV-1 envelopes from virions that persist in plasma on antiretroviral therapy show reduced susceptibility to autologous immunoglobulins and variable sensitivity to broadly neutralizing monoclonal antibodies

**DOI:** 10.1101/2025.05.22.655427

**Authors:** Savrina Manhas, Joseph P. Brooker, Cory Shetler, Kerri J. Penrose, Divya S. Jaiswal, Xiaojie Chu, Wei Li, Mary F. Kearney, John W. Mellors, Elias K. Halvas

## Abstract

Despite adherence to antiretroviral therapy (ART), low levels of plasma virus persist in most individuals with HIV-1. In a small subset of these individuals, this non-suppressible viremia is detected by routine clinical testing (plasma HIV-1 RNA >20 copies/ml) and has been shown to be of clonal cellular origin. The mechanisms by which these virions escape immune clearance is not defined, but reduced binding and neutralization of their surface envelope protein (Env) by antibodies (Abs) may contribute. To assess this possibility, we measured the monoclonal antibody (mAb) sensitivity of HIV-1 Env in plasma virus from 4 well-characterized individuals on ART with non-suppressible viremia. Thirty-two *env* plasma sequences were used to produce pseudovirus for neutralization assays with both autologous plasma and a panel of 15 recombinant antibodies that targeted different regions of HIV-1 Env protein. We found that autologous plasma had no neutralizing activity against the pseudoviruses consistent with persistence of viremia. In general, the Envs from the non-suppressible plasma virus were also less sensitive to mAb neutralization compared controls: tier 1 (6535), tier 2 (TRO11) or tier 3 (PVO) subtype B *envs*, although variability across Envs was evident. Two Env protein variants from one donor, R-09_A8 and R-09_C2, were less sensitive to VRC01 likely due to an additional N-glycan site at a VRC01 contact site and longer V5 regions. Most of the donor Envs were sensitive to at least two of three V3-glycan mAbs except for variant C-03_A6, which showed reduced sensitivity to all three. The CD4 binding site mAb 3BNC117 and the Gp41-specific mAb 10E8 neutralized pseudoviruses from all donors, indicating the potential for clearance of persistent viremia in these individuals studied.

**Author summary:** In a fraction of individuals on suppressive antiretroviral therapy (ART), episodes of persistent low-level viremia can be observed that is not attributed to ineffective ART and/or non-adherence. Previously in three individuals, we have shown that this viremia results from large HIV-1 infected T cell clones harboring replication-competent proviruses. The factor(s) that allow these infectious virions to be detected and not cleared by the immune system is unknown, suggesting that these virions can evade humoral responses. Using single genome sequencing, we amplified the *env* from plasma derived virions from these individuals, cloned *env* -containing amplicons into an expression vector to produce pseudo viruses, which we tested against either their autologous contemporaneous autologous immunoglobulins (Ig) or a panel of 15 recombinant monoclonal antibodies. The results revealed that these pseudovirus were not neutralized by their autologous Igs and exhibited complex pseudovirus-specific susceptibility profiles for the monoclonal antibodies they were tested against. Collectively, our findings suggest that despite resistance to autologous Ig, likely combinations of monoclonal antibodies will be needed to clear this persistent viremia.

## Introduction

Antiretroviral therapy (ART) blocks HIV-1 replication and disease progression (Perelson), (Palella) but does not cure HIV-1 infection because of the long-term persistence of infected cells carrying replication-competent proviruses that cause rapid viral rebound after cessation of ART (de Jong) (Finizi). It is widely acknowledged that elimination or immune control of this pool of infected cells is a major challenge for HIV-1 cure strategies. Despite adherence to ART, residual viremia persists in most individuals. In a small subset of individuals, this non-suppressible viremia is detected by routine clinical testing (plasma HIV-1 RNA >20 copies/ml) and has been shown to be of clonal cellular origin (Halvas 2020). The mechanisms whereby the cellular clones and the virions they produce escape immune clearance are not defined.

One potential mechanism of escape is resistance to humoral immune responses. The target of HIV-1 neutralizing Abs are the envelope glycoprotein (Env) spikes on the surface of virions, composed of three gp120-gp41 dimers (Wyatt). Although most individuals develop HIV-1 Abs during natural infection, only a small percent develop potent antibody responses capable of neutralizing a broad range of variants; such Abs are known as broadly neutralizing Abs (bNAbs) (Simek). HIV-1 bNAbs can be categorized based on the epitopes they recognize such as the CD4 binding site (CD4bs), V1V2 apex, V3-glycan, gp120-gp41 interface, and gp41 membrane-proximal region (MPER) (Burton). Unfortunately, bNAbs have been difficult to induce through vaccination due to their unique features such as a high degree of somatic hypermutation and a relatively long heavy chain complementarity determining region 3 (HCDR3) (Burton).

By contrast, many recombinant bNAbs have been produced and their therapeutic administration has shown promising results in reducing HIV-1 replication (Julg-19, Hsu). Potential pitfalls of using bNAb therapy to control HIV-1 replication, clear virions or eliminate virus-infected cells expressing Env, is the pre-existence of virions that have reduced sensitivity to bNAbs or the evolution of bNAb resistant Env escape mutants (Caskey-17).

In the current study, we focused on both autologous immunoglobulin (Ig) and the mAb sensitivity of Envs derived from plasma virions obtained from 4 individuals with clinically detected, non-suppressible viremia on ART (Halvas). Plasma virus from 3 of the 4 individuals represents the expressed, replication-competent reservoir (Halvas). The neutralization sensitivity of these Envs expressed on pseudoviruses was tested against both autologous Ig and a panel of 15 bNAbs (b12, VRC01, 3BNC117, LSEVh-LS, PGT-121, 10-1074, PGT-128, PG9, PG16, PGT-145, 2F5, 4E10, 10E8, PGT151, N6/PGDM1400x10E8) targeting multiple sites on Env. The sensitivity of the donor Envs to neutralization varied widely, even within donors and across the different classes of bNAbs tested.

## Results

### Study participants and clinical characteristics

Four individuals with HIV-1 infection for a mean of 21 years (range, 10-24 years) and a mean of 14 years on long-term ART (range, 9-19 years) with periods of non-suppressible viremia were studied with the characteristics of the donors provided (Table 1). The mean age of the donors was 60.5 years (range, 43-73 years) and the mean years of clinically detected viremia (>20 copies/ml) was 4.9 years (range, 2.1-7.4 years).

**Table 1.**
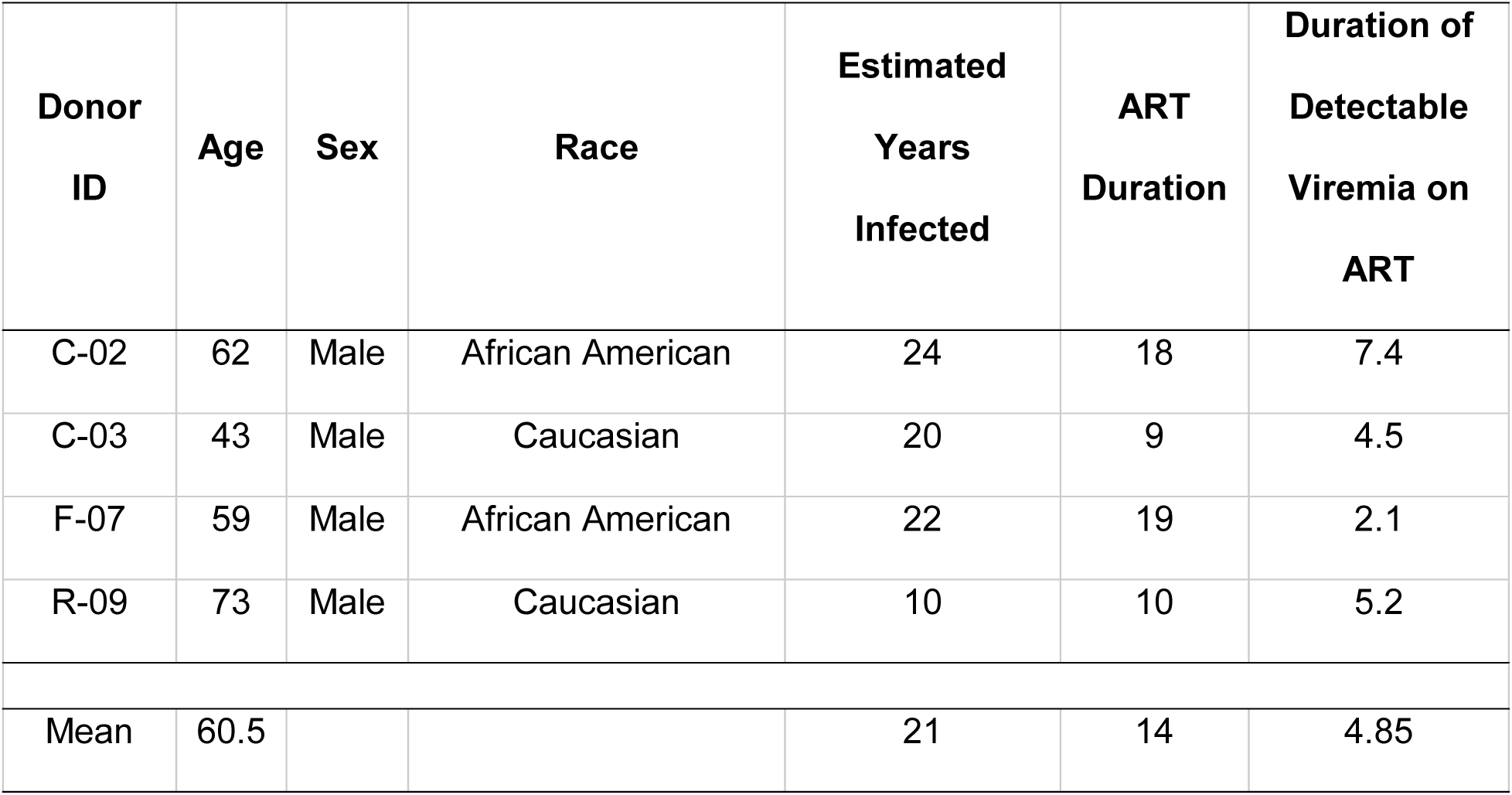
Characteristics of donors with non-suppressible viremia on antiretroviral therapy.

### Single genome amplification and sequencing of HIV-1 *env* from plasma viral RNA

Median plasma HIV-1 viral RNA at time of referral was 18.6 copies/mL (range, 9.2 to 121.7 copies/mL), as determined by the single copy assay (SCA) targeting *pol* (Cillo). Viral RNA was extracted for cDNA synthesis and a nested touchdown PCR approach was used for HIV-1 *env* single genome sequencing (SGS) (Palmer). The median estimated total copies of HIV-1 RNA in the plasma volume tested as tested by SCA was 34 copies (range, 13 to 75 copies) and the median recovery of full-length *env* sequences was 27% (range, 6.7% to 69%) (Table 2).

**Table 2.**
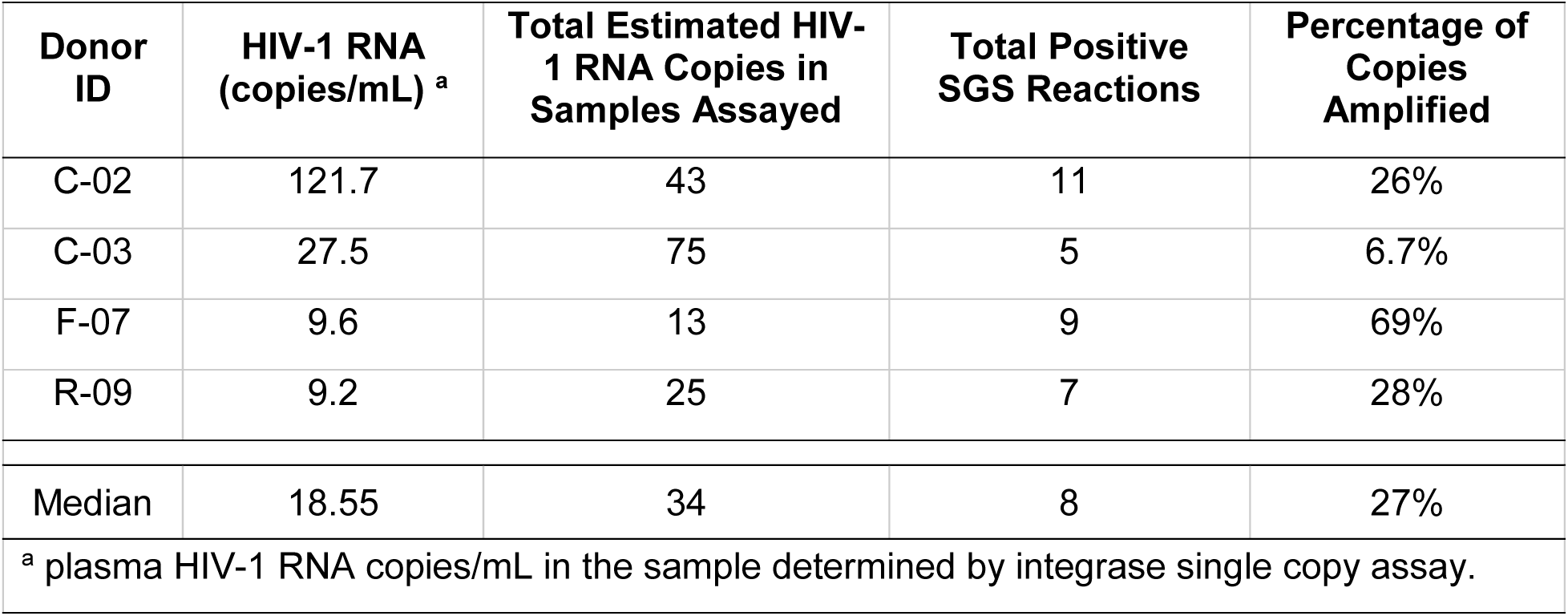
Single genome sequencing (SGS) efficiencies of plasma-derived HIV-1 envelopes from donors with non-suppressible viremia on ART.

We generated and sequenced, using the Illumina MiSeq platform, a total of 32 HIV-1 *env* amplicons across four donors. Phylogenetic analyses of the HIV-1 *env* sequences revealed distinct clustering with no overlap between the four donors (Fig 1) (Kumar). Each donor had identical *env* sequences, including 11 of 11, 4 of 9, and 2 of 5 sequences for donors C-02, F-07, and C-03, respectively. Donor R-09 had two distinct clusters of identical *env* sequences (3 versus 4 sequences), with the clusters differing at 52 nucleotide positions in *env*. All the identical sequences from donor C-02, two from donor C-03, and one of the *env* sequence clusters from donor R-09, matched clones of cells with replication-competent proviruses that were previously report (Halvas). Some of donors (C-03 and F-07) also had plasma *env* sequences that were divergent from these identical sequences by a single to 334 nucleotides.

**Fig 1.**
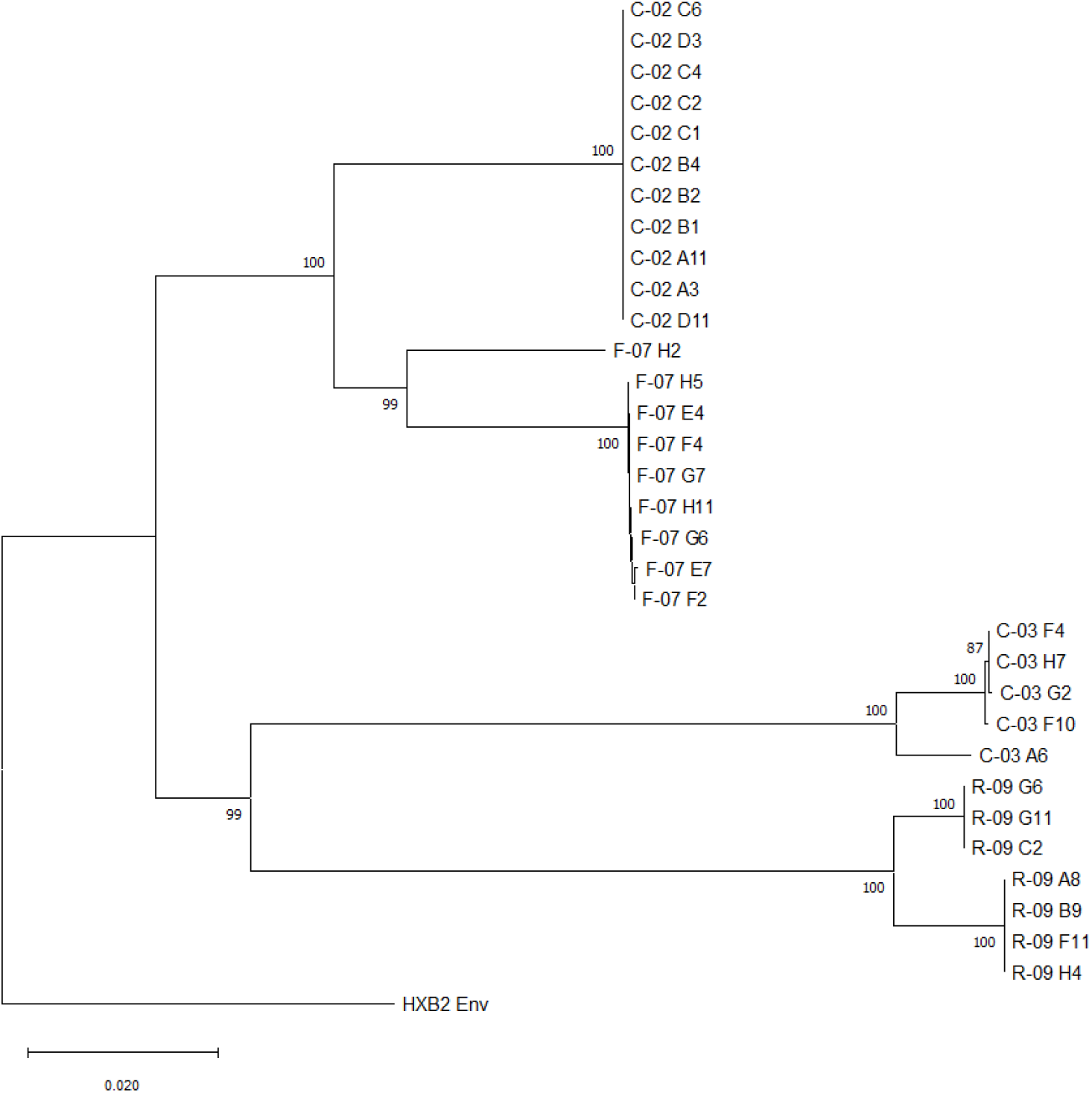
Neighbor-joining tree of HIV-1 env sequences amplified from plasma virus from donors C-02, C-03, F-07, and R-09. The evolutionary history was inferred using the Neighbor-Joining method. The optimal tree is shown. The percentage of replicate trees in which the associated taxa clustered together in the bootstrap test (1000 replicates) are shown next to the branches. The tree is drawn to scale, with branch lengths in the same units as those of the evolutionary distances used to infer the phylogenetic tree. The evolutionary distances were computed using the p-distance method and are in the units of the number of base differences per site. This analysis involved 33 nucleotide sequences. Codon positions included were 1st+2nd+3rd+Noncoding. All ambiguous positions were removed for each sequence pair (pairwise deletion option). There were a total of 2682 positions in the final dataset. Evolutionary analyses were conducted in MEGA X (13) (Kumar-2018). The tree was rooted to the HXB2 reference sequence.

### Neutralization of pseudoviruses with HIV-1 envelopes from donor plasma

To evaluate the activity of autologous Ig and specific monoclonal antibodies against isolated donor derived envelopes, we ligated unique HIV-1 *env* amplicons from each donor which were generated by SGS into a mammalian expression vector and co-transfected 293T cells along with an HIV-1 luciferase reporter plasmid to generate pseudoviruses (PSVs). Neutralization assays were performed by incubating PSVs with dilutions of either donor-derived autologous Ig or a panel of monoclonal antibodies (Abs) followed by addition to TZM-bl cells and subsequent assessment of luciferase activity 72h later. Negative controls to ascertain signal from background included incubating PSVs in the absence of either autologous Ig or the panel of 15 monoclonal Abs used to determine neutralization sensitivities.

### Reduced susceptibility of plasma-derived pseudoviruses to autologous IgG

Previous studies in persons with HIV (PWH) and on suppressive ART with <40c/mL have reported variable neutralization susceptibilities to both heterologous and autologous viruses interrogated by autologous IgG (Gartner, Wilson). Yet others, also in PWH on suppressive ART have observed that >80% of viruses are inhibited by autologous IgG in 40% of the participants tested using a modified viral outgrowth assay and measuring differences in infectious units per million cells (Bertagnolli). We hypothesized that our PSVs with HIV-1 Envs derived from plasma could at the least exhibit variable neutralization sensitivities to contemporaneous autologous Igs as previously reported (Gartner, Wilson). In contrast, due to the non-suppressible nature of our participants and the likelihood that these PSVs are not readily neutralized and cleared by the immune system, we could also observe no effect from the autologous Igs.

To gain insight on how these viruses can evade the immune system, we tested our PSVs containing plasma-derived Envs against their respective autologous Igs. The primary concentration of purified protein was set at 500ug/mL as previously described (Esmaeilzadeh). This concentration was 10-20× the primary concentration reported in previous studies (Gartner, Wilson, Bertagnolli) and 200× the concentration used with the panel of monoclonal antibodies in this study. All PSVs and controls (i.e. 6535, TRO.11, and PVO) exhibited reduced sensitivity to neutralization by their respective autologous Igs (range; 500ug/mL to 0.009ug/mL) (Figs 2A-2D). TRO.11 and PVO were also tested against 10-1074 as an assay run control showing inhibition of the control PSVs (Fig 2E). The presence of purified total Ig was confirmed by Sodium Dodecyl Sulfate-Polyacrylamide Gel Electrophoresis (SDS-PAGE) and the concentrations of purified IgG1-G4, IgA, and IgM proteins used in the neutralization assays were determined by quantitative enzyme-linked immunosorbent assays (ELISA) (S1 Fig) and (S1 Table). The median concentration of purified IgG1-G4 among the four donors was 1.94mg/mL (range, 1.489-2.596mg/mL) with IgG making up a median of ∼18% of the Igs in the purified samples.

**Fig 2.**
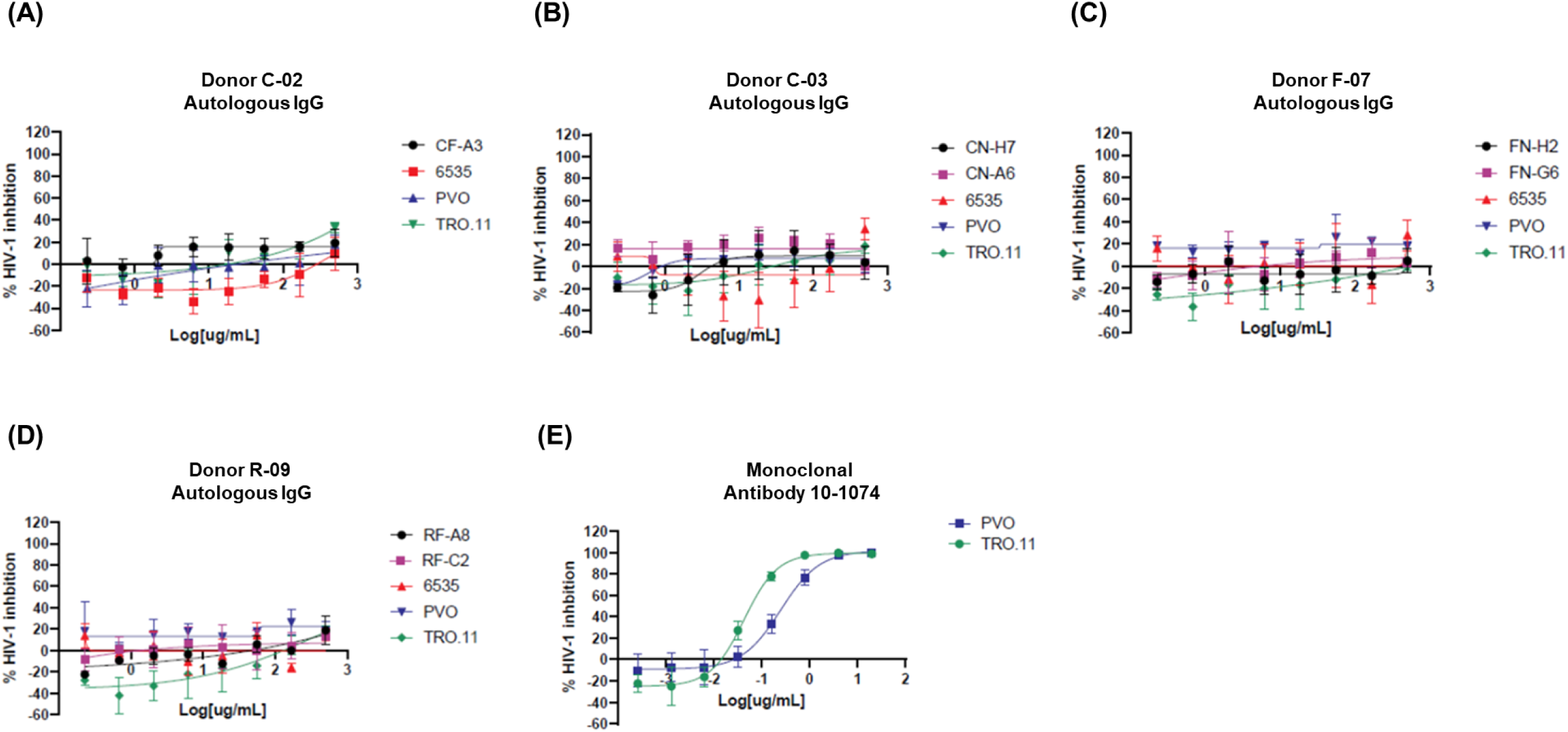
Neutralization sensitivities for participant-derived autologous IgGs. Donors (A) C-02, (B) C-03, (C) F-07, and (D) R-09 HIV-1 envelopes were expressed as pseudovirions and incubated with their respective autologous IgGs for 1 h and then added to TZM-bl target cells. Luciferase was measured after 3 days. Percentage neutralization was calculated relative to cell-only and virus-only controls. The subtype B pseudoviruses 6535, TRO.11, and PVO were used in parallel as neutralization controls. (E) Control pseudoviruses TRO.11 and PVO were also tested against monoclonal antibody 10-1074 as an assay run control. Neutralization curves fitted by GraphPad PRISM v10.0.0.

### Neutralization of pseudoviruses with HIV-1 envelopes by monoclonal antibodies

Given that an in-vivo concentration of ∼200-fold above the neutralization IC50 measured in vitro in TZM-bl cells is generally needed to ensure 100% protection against viral challenge in passive transfer non-human primate (NHP) studies, Abs with an IC50 of <1 µg/ml in the TZM-bl neutralization assays are considered to be most realistic choices for prophylaxis (Sok). As such, we tested our Abs at 1:5 serial dilutions starting at 2.5 µg/ml. Antibody resistance to neutralization was defined as IC50 at >2.50 µg/ml.

Alongside the donor PSVs, we also tested the neutralization of the Ab panel against PSVs generated from the tier 1b subtype B Env 6535 (S1-S6 Figs) (Table 3) to ensure assay accuracy and reproducibility, as the neutralization susceptibility of this HIV-1 Env has been well characterized (Li-05). In general, each of the donor PSVs tested had a unique neutralization profile, even those isolated from the same donor (Table 3). PSV R-09_C2 was the least sensitive to neutralization being less sensitive to 11 of the 15 Abs tested, whereas PSV F-07_H2 was the most susceptible, showing insensitivity to only 2 of 15 Abs.

**Table 3.**
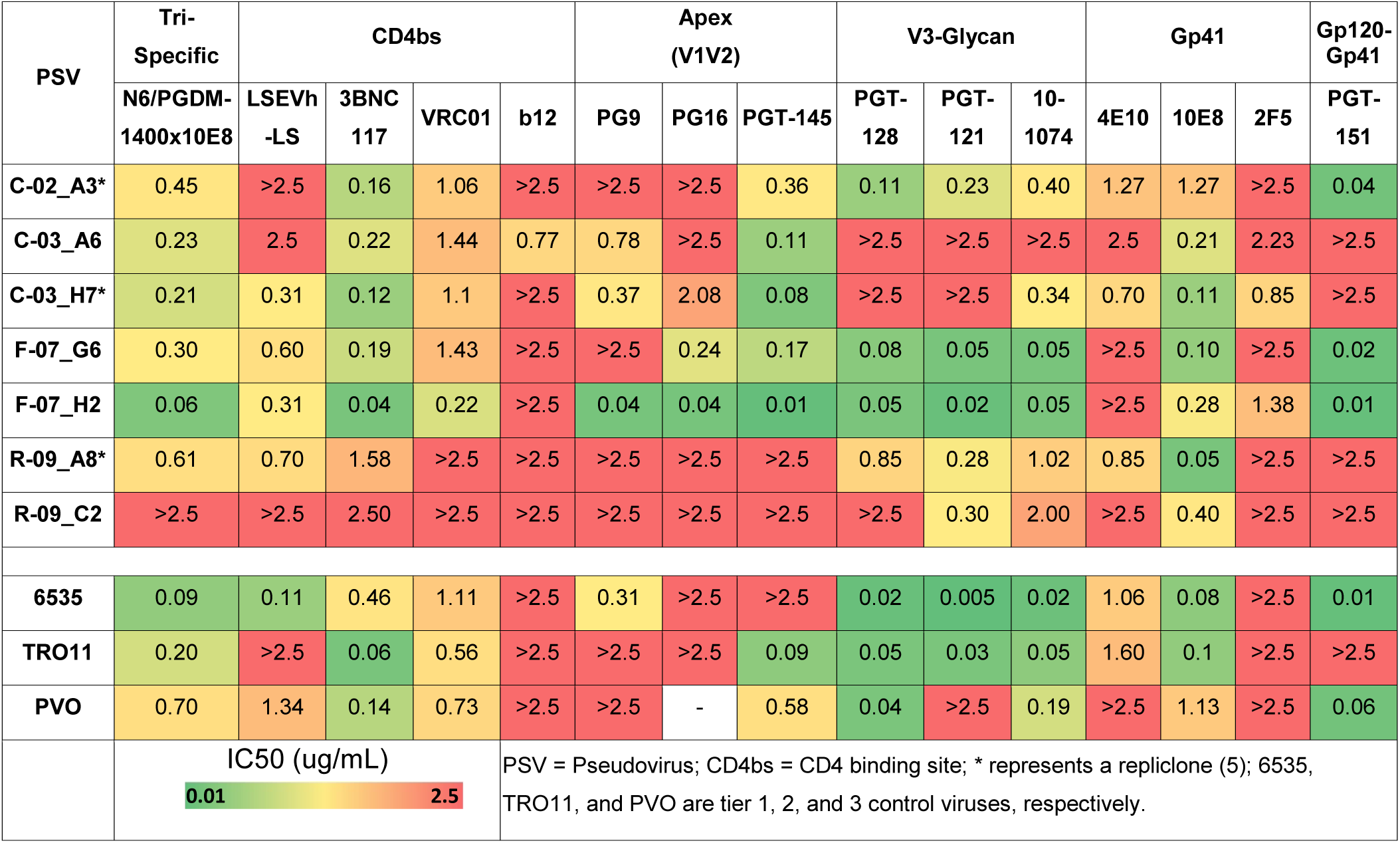
Neutralization sensitivity of plasma-derived HIV-1 envelopes from donors with residual viremia on ART.

### Susceptibility by monoclonal antibody binding to CD4 binding site of gp120

The first epitope we probed was the functionally conserved CD4 binding site (CD4bs) on gp120 that binds to the host cell receptor CD4 (Kwong). Given that this initial step is essential for entry, the CD4bs is functionally conserved on Env and vulnerable to Ab neutralization. We tested three mAbs (VRC01, 3BNC117 and b12) and one fusion protein (LSEVh-LS) directed to the CD4bs. The trend in neutralization of PSVs by mAbs targeting the CD4bs from most active to least was 3BNC117 > VRC01> LSEVh-LS >b12, with 7 of 8 PSVs tested exhibiting reduced sensitivity (>2.5 µg/ml) to b12 (S1 Fig) (Table 3). Of note, PSV RF_C2 showed resistance to neutralization by all CD4bs mAbs tested except for 3BNC117 (IC50 = 2.50 µg/ml) (S1 Fig) (Table 3). Furthermore, all PSVs tested here were generally less sensitive to neutralization by CD4bs mAbs than the 6535 tier 1 control PSV (S2 Fig).

The fusion protein LSEVh-LS is composed of 4 engineered single extracellular domain CD4 molecules (mD1.22) and 2 m36.4 antibody domains that target the co-receptor binding site on gp120 (Chen-16). Additionally, a heavy chain constant domain, kappa light-chain constant domain and an IgG Fc domain are fused to the mD.122 and m36.4 molecules to generate the hexavalent binding protein LSEVh-LS (Chen-16). PSV R-09_C2 and C-02_A3 were less sensitive to LSEVh-LS neutralization, whereas C-03_A6 was 50% neutralized at 2.5 µg/ml. which was the highest LSEVh-LS concentration tested (S2 Fig) (Table 3). In examining the sequences of the *envs* with resistance or reduce susceptibility, we observed no obvious signature sequence, specifically at the amino acid positions postulated to interact with CD4, which could account for resistance to neutralization by LSEVh-LS (Wu-09, Zhou-19) (Fig 3).

**Fig 3.**
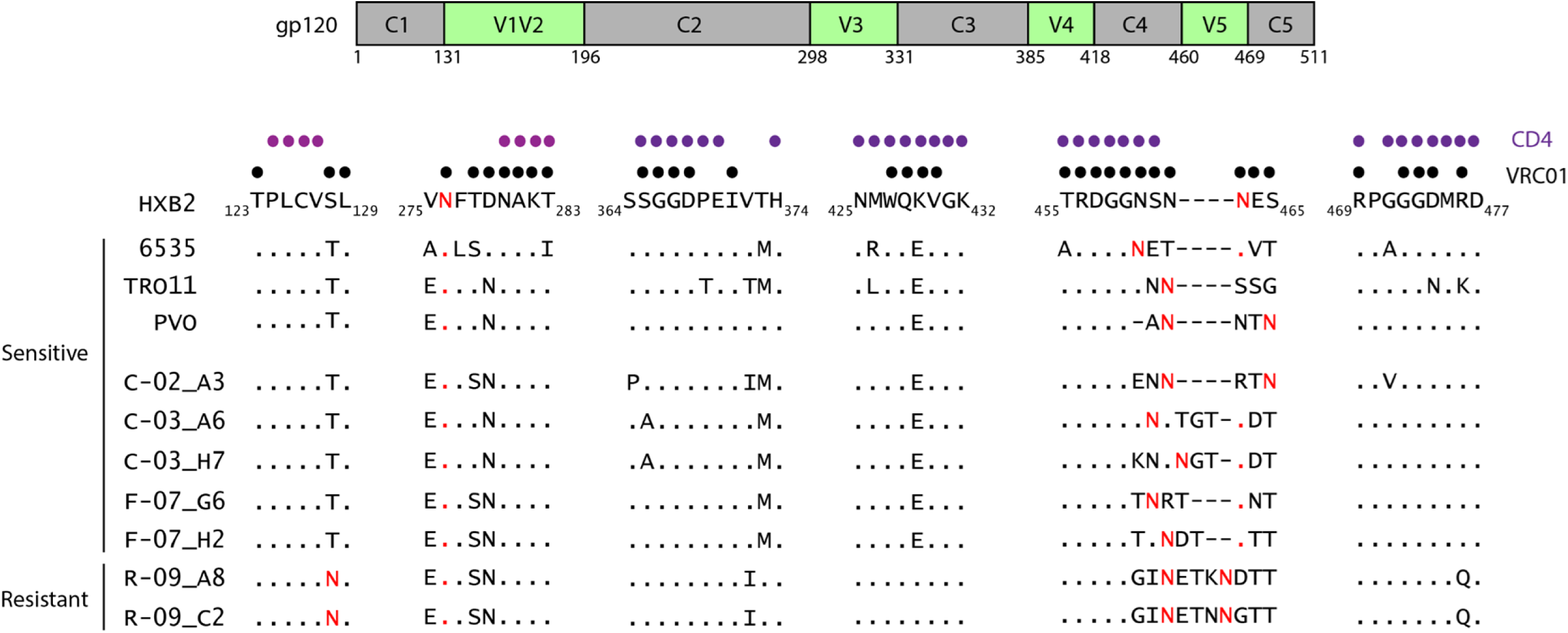
Additional potential N-linked glycosylation sites (PNGS) and changes in V5 affect neutralization by VRC01. A schematic of gp120 is shown at the top with the sequence variable regions V1 to V5 in green and the sequence conserved regions C1 to C5 in grey; regions are numbered according to HXB2. The protein sequence of the CD4bs and VRC01 epitope was analyzed for donor Env sequences (C-02, C-03, F-07, and R-09) along with the following subtype B Envs controls: 6535 (tier 1b), TRO11 (tier 2) and PVO (tier 3). Previously defined CD4 contact residues (16,18,19) (Kwong, Wu-09, Zhou-19) are denoted by purple circles and VRC01 contact residues (57) (Zhou-19) by black circles. Asparagine residues (N) highlighted in red are PNGS with the motif N-X-T/S where X denotes any amino acid residue. Small black dots represent homology to the HXB2 reference sequence and small red dots represent PNGS residues conserved in HXB2. Dashes denote regions that tolerate amino acid insertions.

### Susceptibility by monoclonal antibody binding to V3 binding site of gp120

The V3 region on gp120 is required for host cell co-receptor binding. mAbs targeting the V3 binding site primarily target the V3 glycans N295, N301, and N332, along with other V3 residues (Burton-16). We assessed the neutralizing activity of the V3-glycan Abs PGT-121, PGT-128 and 10-1074 against our panel of PSVs. Results revealed PSVs to be most sensitive to 10-1074 > PGT-121 >PGT-128, and all donor PSVs were less sensitive than the 6535 and TRO11 PSV controls (S3 Fig) (Table 3). PSV C-03_A6 was less sensitive to all three Abs tested. PSV C-03_H7 isolated from the same donor was less sensitive to PGT-121 and PGT-128 but was sensitive to 10-1074 (0.34 µg/ml) (S3 Fig) (Table 3).

### Susceptibility by monoclonal antibody binding to V1/V2 binding site of gp120

A previous study revealed that the V1V2 and V3 loops from three gp120-g41 protomers of the Env trimer converge at the membrane-distal portion of the spike (Kwon) and that the V1V2 region on gp120 is vulnerable to neutralizing Abs (Doria-Rose). We tested the neutralizing activity of V1/V2 targeting Abs PG9, PG16, and PG145. In general, PSVs tested were more sensitive to neutralization by PGT-145 than PG9 and PG16; however, sensitivity varied widely by PSV. For example, both PSVs from donor R-09 showed resistance to neutralization to all three V1V2 apex Abs (>2.5 µg/ml). By contrast, PSV F-07_ H2 was sensitive to all three Abs (IC50s ranging from 0.01-0.04 µg/ml) and the remaining PSVs had various degrees of sensitivity (S4 Fig) (Table 3).

### Susceptibility by monoclonal antibody binding to MPER binding site of gp41

The gp41 MPER is critical for viral and host cell membrane fusion and is a vulnerable site for Ab binding and neutralization (Montero). We tested the neutralizing activity of three gp41 MPER targeting Abs: 4E10, 10E8 and 2F5 against the donor-derived and control PSVs. All PSVs displayed sensitivity to 10e08 and various degrees of resistance to neutralization by 4E10 and 2F5 (S5 Fig) (Table 3). None of the PSVs tested were resistant to neutralization by all three Gp41 Abs, however four of seven PSVs were less sensitive to neutralization by 2F5 (S5 Fig) (Table 3).

### Susceptibility by monoclonal antibody binding to gp120-41 interface

Previous studies have shown that PGT151 binds a quaternary epitope present on the HIV-1 Env trimer at the interface of gp120-gp41 (Blattner). Interestingly, sensitivity or resistance to neutralization of donor-derived PSVs by PGT151 appeared to be donor specific (S6 Fig) (Table 3). The PSVs from donors R-09 and C-03 exhibited diminished sensitivity to neutralization, whereas those from the remaining donors were sensitive (IC50 range, 0.01 to 0.04 µg/ml) (S6 Fig) (Table 3).

### Susceptibility by Trispecific monoclonal antibody

We tested all seven PSVs against the tri-specific mAb N6/PGDM1400x10E8, which targets CD4bs, V1V2 apex, and MPER binding sites (Xu). All PSVs showed neutralization sensitivity to the trispecific antibody except PSV R-09_C2, which was less sensitive (S7 Fig) (Table 3). This finding of resistance was somewhat expected given the degree of reduced susceptibility of the R-09_C2 to CD4bs, V1V2, and MPER Abs.

### Variable region lengths and number of potential N-linked glycosylation sites (PNGS) are associated with neutralization resistance

Mutations in *env* allows HIV-1 to evade humoral immune recognition through masking of residues or epitopes that are targets of Ab binding. It has been extensively reported that two mechanisms, either longer variable regions, particularly V1V2 or V3, or increased potential N-linked glycan sites (PNGS), are correlated with neutralization resistance (Ringe, van Gils-11, Doores, Goo). We therefore calculated the lengths of the variable regions (V1V2 through V5) as well as the number of PNGS in the donor HIV-1 *env* sequences we analyzed as PSVs. Donor HIV-1 *env* sequences had mean lengths of 70, 35, 31, and 13 amino acid residues for variable regions V1V2, V3, V4, and V5, respectively, which were variable compared to the controls (Table 4). However, the V1V2 variable loops among the donors exhibited the greatest variability in length (range, 68-85 amino acids) with the most pronounced differences 1 to 17 residues versus 7 to 12 for controls. By contrast, V3, V4 and V5 regions had less pronounced differences in amino acid lengths (range, 0-5 residues) (Table 4).

**Table 4.**
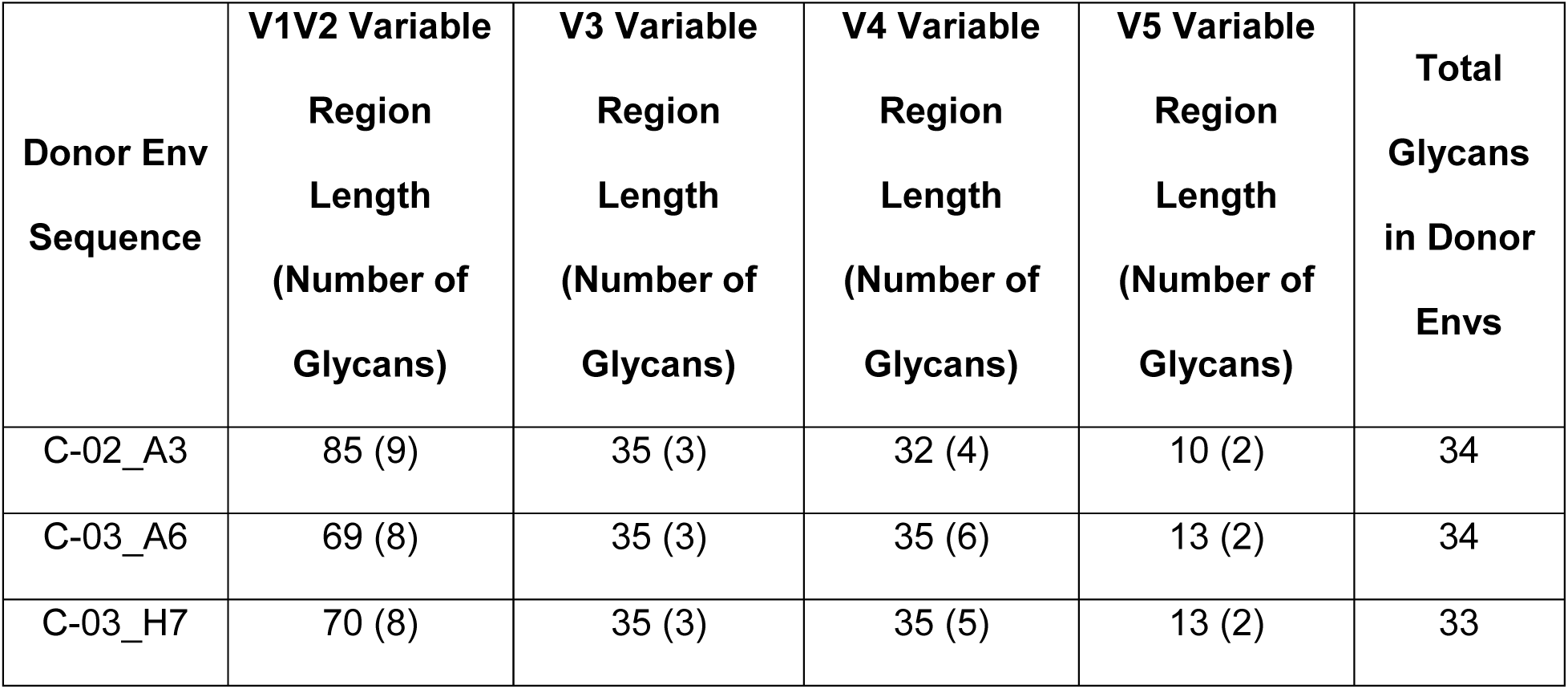
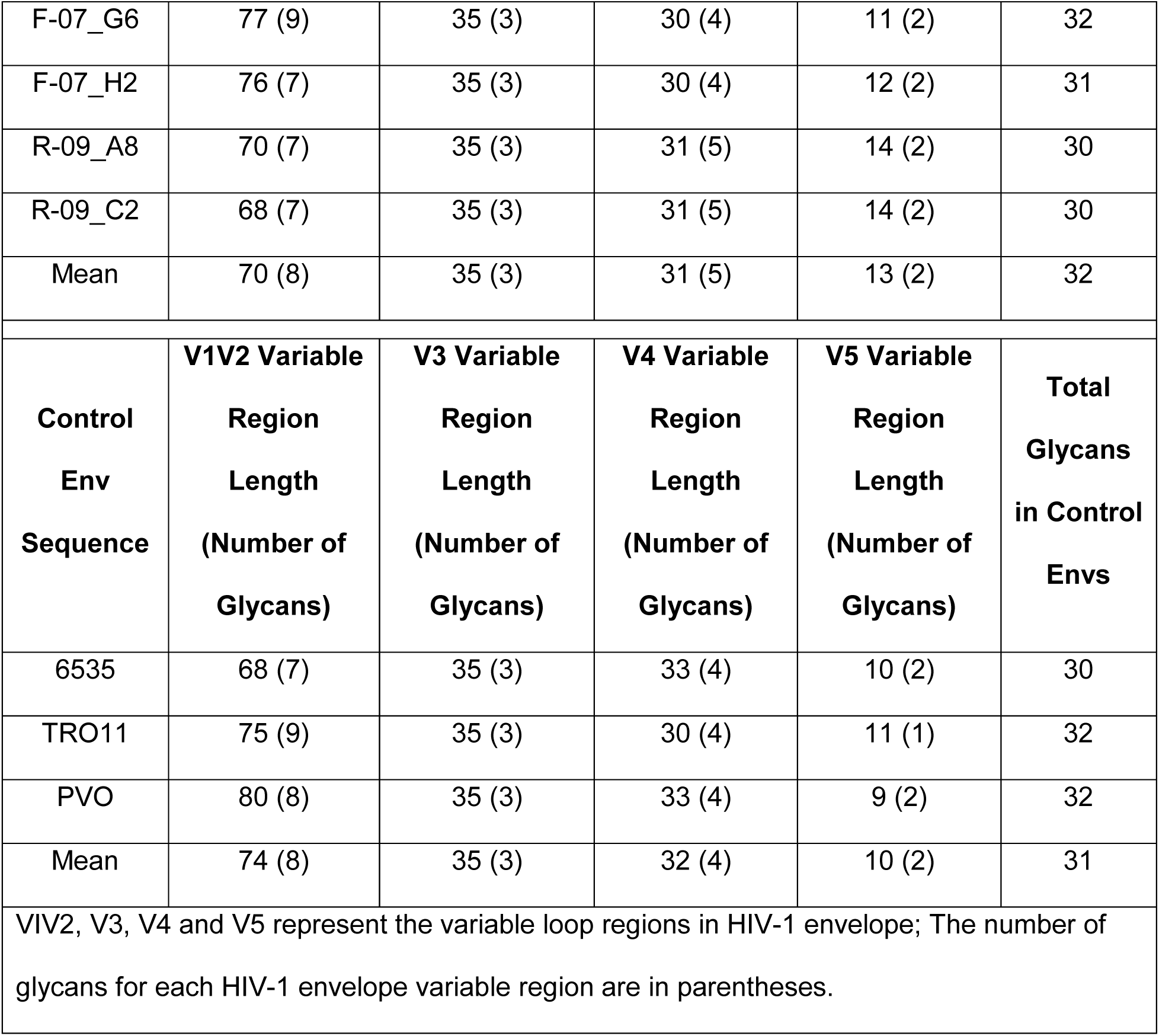
Variable region length and number of glycans in plasma-derived HIV-1 envelopes in donors with persistent viremia on ART.

As for PNGS sites, we found a mean of 32 PNGS in the donor HIV-1 *env* sequences versus the 31 in the controls. The number of PNGSs differed across donor HIV-1 Env by a maximum of 2 PNGS in V1V2 and V4, whereas the number of PNGS was constant in V3 and V5 (Table 4). Although the V1V2 displayed the most dramatic differences in lengths, these did not significantly associate with sensitivity to neutralization for Abs tested (i.e. CD4bs, V3-glycan, or gp120-gp41 interface), nor did the number of PNGS (Data not shown).

### PNGS, substitutions, and longer V5 region could be modulating VRC01- or b12-resistance

Next analyzed were the *env* sequences for mutations in Ab epitopes to characterize the neutralization profiles we observed above (Figs 1 and S2-S7) (Table 3). VRC01 is one of the best-studied CD4bs Ab with considerable neutralization breadth and previously shown to neutralize ∼90% of circulating HIV-1 isolates (Wu-10). Of the donor HIV-1 Envs studied here, variants R-09_A8 and R-09_C2 were both less sensitive to the CD4bs Ab VRC01 (S2 Fig) (Table 3). We analyzed the CD4 and VRC01 contact residues in the donor R-09 HIV-1 *env* sequences to identify residues that could potentially be contributing to VRC01 resistance. In comparison to the VRC01-sensitive HIV-1 *env* sequences, the VRC01-resistant Envs of R-09_A8 and R-09_C2, had i) the addition of a PNGS at position 128, a VRC01 contact residue and ii) the largest number of amino acid additions in V5 (Fig 3) (Table 4).

All donor-derived HIV-1 Envs except variant C-03_A6, exhibited resistance to by Ab b12. Various amino acid positions in gp120 (i.e. H364, P369, and M373) and in gp41 (i.e. T569 and I675) as well as PNGS (i.e. N197 and N301) have been implicated in b12 resistance (Wu-09). All of our HIV-1 Envs retained i) both PNGS at amino acid positions 197 and 301 of gp120, ii) the amino acid substitutions T569 and I675 in gp41, as well as the P369 mutation, whereas positions 364 and 373, were more variable among the donor-derived Envs (S2 Table). Interestingly, donor C-03 HIV-1 Envs exhibited different b12 sensitivities to neutralization (variant C-03_A6; 0.77 µg/mL versus variant C-03_H7; >2.5 µg/mL), despite identical substitutions at these positions (Table 3). Variance analyses revealed that donor C-03 HIV-1 Envs differed at 21 amino acid positions (S3 Table), likely contributing to their variable neutralization susceptibilities.

### Envelope sequences are mostly sensitive to V3-Glycan Abs

As mentioned previously, the V3 region is required for host co-receptor binding typically to CCR5 or CXCR4 of which binding is dependent on the net charge of the V3 region. Sequences from our donors revealed a reduction in the overall net charge of the V3 loop relative to HXB2 (+4 to +5 versus +10 for HXB2 (Fig 3A). Furthermore, V3 is also a target of the V3-glycan neutralizing Abs PGT-121, PGT-128 and 10-1074 and all HIV-1 *env* sequences analyzed contained the presence of the three N-linked glycosylation sites at amino acid positions 295, 301, and 332 as well as the GD/NIR motif (Fig 4A). Although the majority (71%) of our HIV-1 Envs tested were sensitive to the V3-glycan neutralizing Abs, we found that the two from donor C-03 were less sensitive to neutralization by PGT-121 and PGT-128 Abs. However, we observed no obvious differences in the overall net charge or N-linked glycosylation sites with those that were less sensitive versus those that were sensitive (Figs 4A and S3) (Table 3). A comparison of those *env* sequences sensitive to neutralization by PGT-121 and PGT-128, versus those less sensitive, revealed a neutral glutamine residue at position 330 in the Env sequences from donor C-03, while most other Env sequences had a positively charged histidine (Fig 4A). Despite, identical V3 *env* sequences from donor C-03, variant C-03_A6 also exhibited resistance to 10-1074, suggesting residues outside of the epitope or Env conformation can mediate Ab neutralization sensitivity. In addition, R-09_C2 Env displayed resistance to neutralization by PGT-128, but no differences in charge or amino acids in V3 could explain the resistance (Fig 4A).

**Fig 4.**
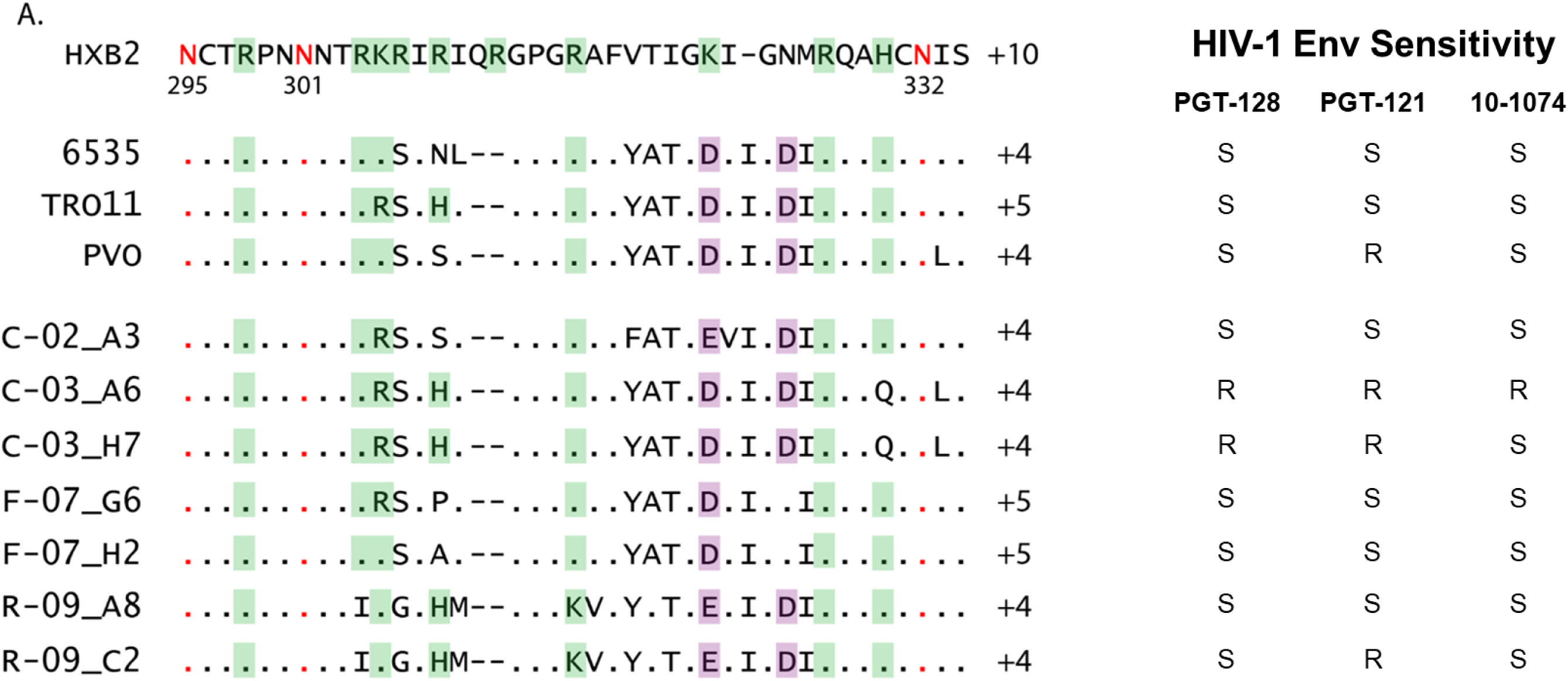
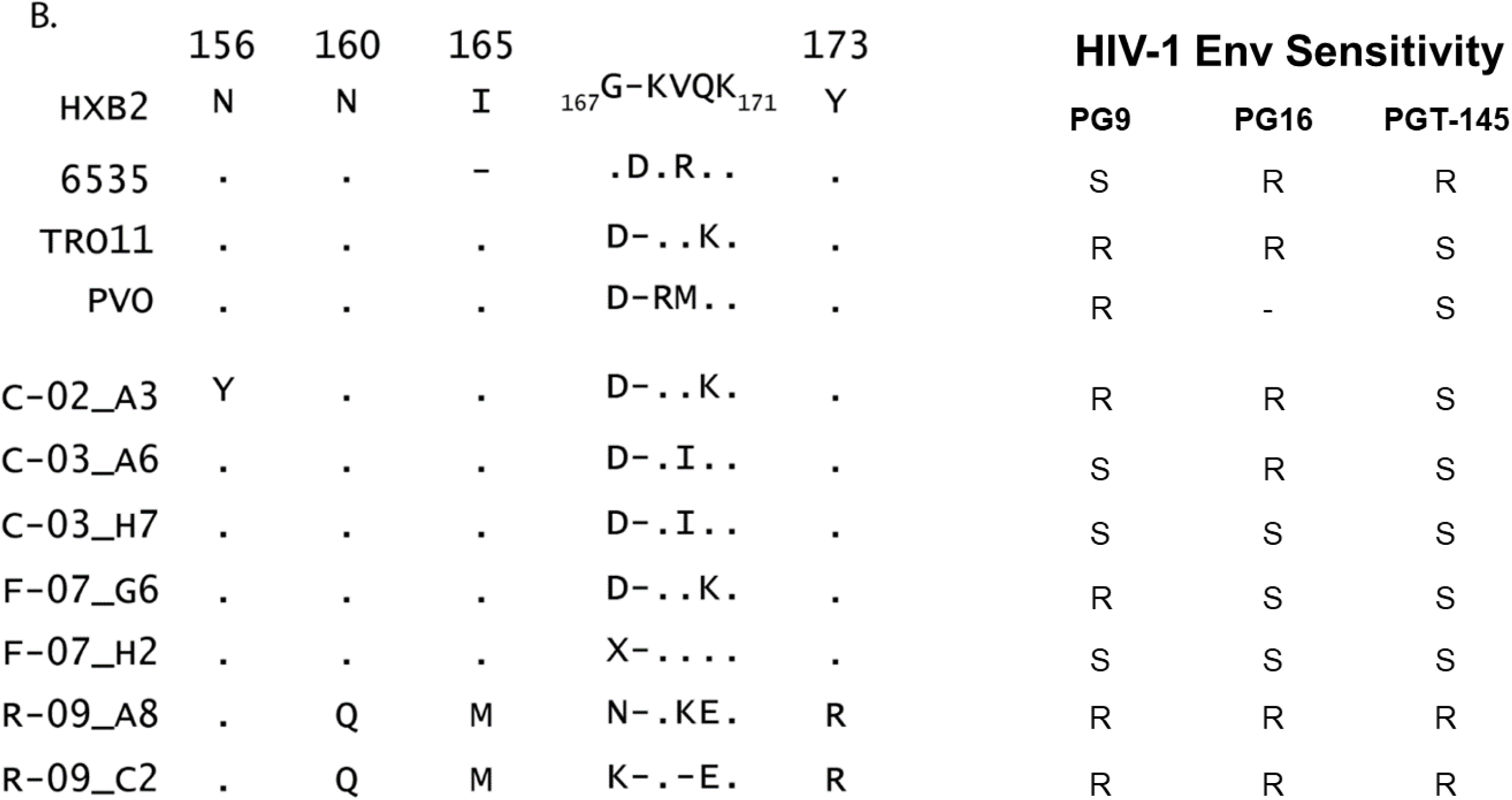
Env sequences lacking N160 render them susceptible to V1V2 apex bNAbs. (A) The V3 sequences of envelopes from C-02, C-03, F-07, and R-09 are shown along with the V3 sequences for envelopes 6535, TRO11 and PVO. The V3 PNGS at N295, N301 and N332 are highlighted in red and numbered. Residues highlighted in green (R, K, H) are positively charged whereas those highlighted in purple are negatively charged (D, E). (B) V1V2 residues that were previously determined as PG9 and PG9-related Ab contacts (21,27) (Doria-Rose, Ringe) are shown for C-02, C-03, F-07, R-09, 6535, TRO11 and PVO envelopes sequences. Residues are numbered according to the HXB2 reference sequence which is shown at the top. (A, B) Black dots represent homologous residues to the HXB2 reference sequence and a dash denotes an area that tolerates a residue insertion. Sensitivities of envelopes to Apex V1V2 and V3-glycan antibodies are listed to the right in panels A and B. S represents envelopes that are sensitive/susceptible and R represents those that are resistant (>2.5mg/mL) to antibodies tested.

### Disruption of N160 PNGS and substitutions in V1V2 results in resistance to neutralization by V1V2 Abs

The V1V2 region on gp120 is not required for viral entry, but is critical for immune evasion as removal of V1V2 renders viruses extremely sensitive to Ab neutralization (Stamatatos). The V1V2 regions of the donor sequences were variable in length (range, 68 to 85 residues), carried 7 to 9 PNGS, which encompasses ∼10% of V1V2 residues, and a number of substitutions spanning residues 156 through 176 (Fig 4B) (Table 4). We analyzed key V1V2 residues in this region implicated in PG9, PG16 and PGT-145 neutralization for addition or subtraction of PNGS or substitutions that would affect the charged interaction between HIV-1 Env and Ab (Doria-Rose). We found that R-09_A8 and R-09_C2 Envs, which were less sensitive to PG9, PG16, and PG-145 neutralization (S4 Fig) (Table 3), lacked the glycan at position 160, gained a PNGS in amino acid position 128 (associated with PG9-resistance), and contained a number of substitutions at other key amino acid positions (i.e. 173) (Figs 3 and 4B) (Doria-Rose). Furthermore, C-02_A3 HIV-1 Env, which also showed neutralization resistance to PG9 and PG16 (S4 Fig) (Table 3), carried a tyrosine residue at position 156, abolishing the glycan site in the V1V2 variable region previously associated with PG9 binding (Fig 4B) (Doria-Rose). By contrast, we observed mixed susceptibilities of neutralization by PG9, PG16, and PG145 for donors C-03 and F-07 (Table 3), despite the preservation of the key PNGS at positions 156 and 160 (Fig 4B). Previous studies have also implicated the overall electrostatic charge of V2 residues 163-176 with alteration of PG9 and PG16 susceptibility, with greater charge associated with neutralization sensitivity (Doria-Rose, Ringe). An inspection of this region revealed no clear pattern of substitution that would account for their variable sensitivities (S4 Table).

### Residues outside of Gp41 or MPER accessibility to Ab affect Gp41-specific neutralization

The MPER of gp41 is the binding site for mAbs 10E8, 4E10 and 2F5, with residues 666, 672, and 673 implicated in neutralization sensitivity/resistance (Wibmer). We therefore analyzed MPER sequences of the donor HIV-1 Env sequences at these key amino acid positions, but did not observe any of the common substitutions that are linked to changes in neutralization (Fig 5). Furthermore, across donors, we saw no clear association between the diverse range of sensitivity to neutralization by the gp41-specific Abs and polymorphisms in the ectodomain of Env sequences (Fig 5) (Table 3). This lack of association was also evident within donors (C-03_A6 versus C-03_H7 or R-09_ A8 versus R-09_C2) with identical ectodomain sequences, but variable profiles of neutralization for Abs 4E10 (Fig 5) (Table 3). An additional finding was the absence of any known substitutions (i.e. leucine, arginine, or glutamine) at HIV-1 Env position 673, which are associated with resistance to antibodies that target the MPER of the ectodomain (Fig 5).

**Fig 5.**
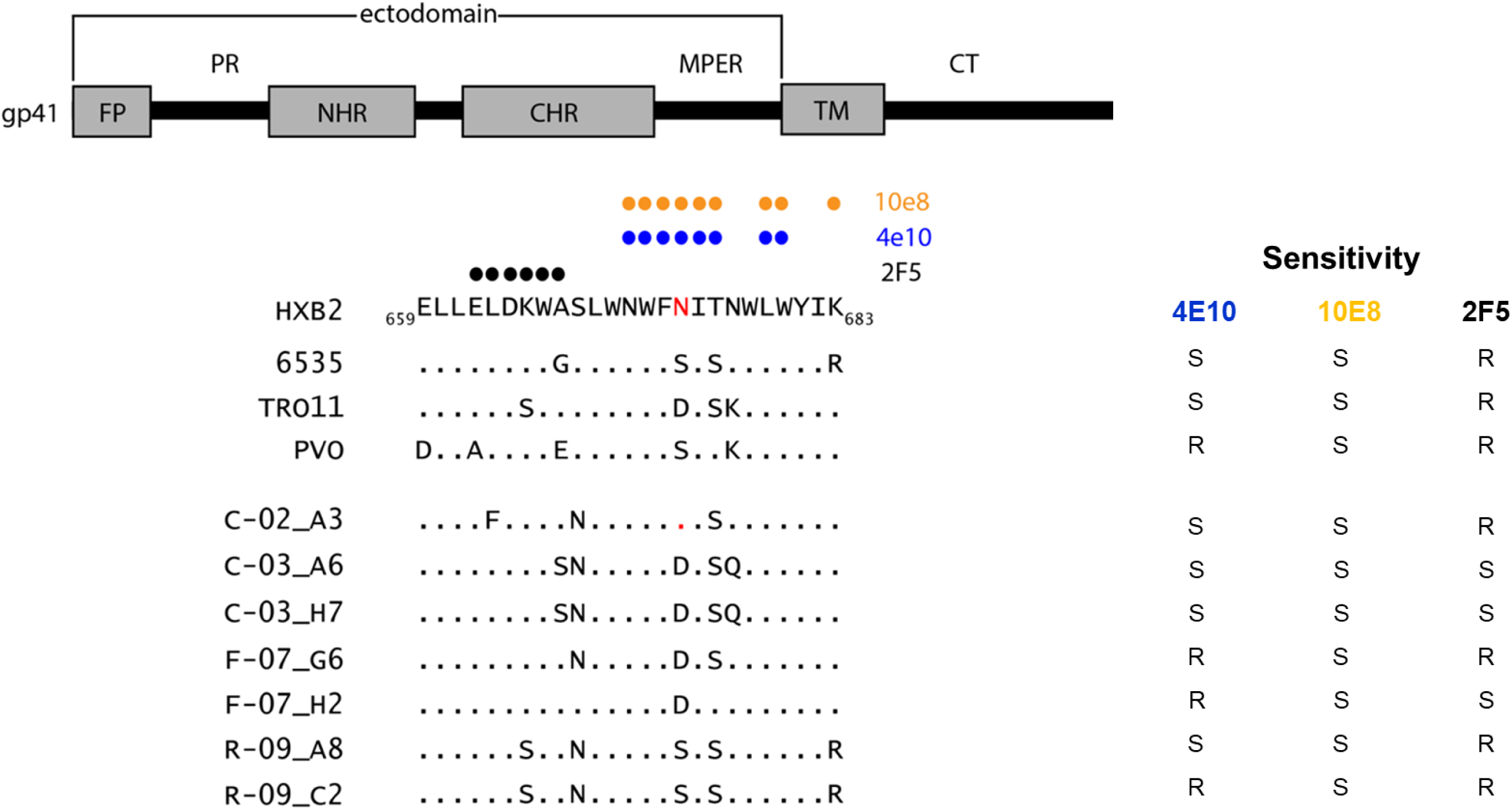
Minor changes in the MPER region of gp41 do not drastically affect gp41-specific neutralization. A schematic of gp41 adapted from (22) (Montero) is shown at the top with the 3 regions (ectodomain, transmembrane (TM) domain and cytoplasmic tail (CT)). The fusion peptide (FP), N-terminal heptad repeat (NHR), C-terminal heptad repeat (CHR) and TM regions are depicted in grey boxes. The polar region (PR), membrane-proximal external region (MPER) and CT are shown in black. Donor envelope MPER sequences are shown along with reference sequences from 6535, TRO11 and PVO. MPER residues that contact 10E8, 4E10 and 2F5 are denoted with orange, blue and black circles. respectively. Black dots represent homology to the HIV-1 HXB2 reference sequence. Sensitivities of envelopes to gp41-specific antibodies are listed to the right. S represents envelopes that are sensitive/susceptible and R represents those that are resistant (>2.5mg/mL) to antibodies tested.

#### Substitutions in Gp41 modulate the neutralizing activity of PGT151

The PGT151 epitope includes inter- and intraprotomer contacts with the Env trimer and is dependent on a subset of glycans, specifically N611 and N637, which are required for PGT151 binding or decreased neutralization (Blattner, van Gils-16). We found that resistance to PGT151 neutralization was donor specific and observed only in donors C-03 and R-09, with those Envs from donor C-03 showing less pronounced resistance (S6 Fig) (Table 3). All four Envs from these donors lacked previously described substitutions at key amino acid positions (i.e. 512, 514, 611, and 637) and therefore, did not provide clear insight on their PGT151 resistance profiles (Fig 6). However, Envs from donor R-09 did lack the glycine residue at position 514, whereas in the Envs from donor C-03, the glycine was present (Figs 6 and S6) (Table 3).

**Fig 6.**
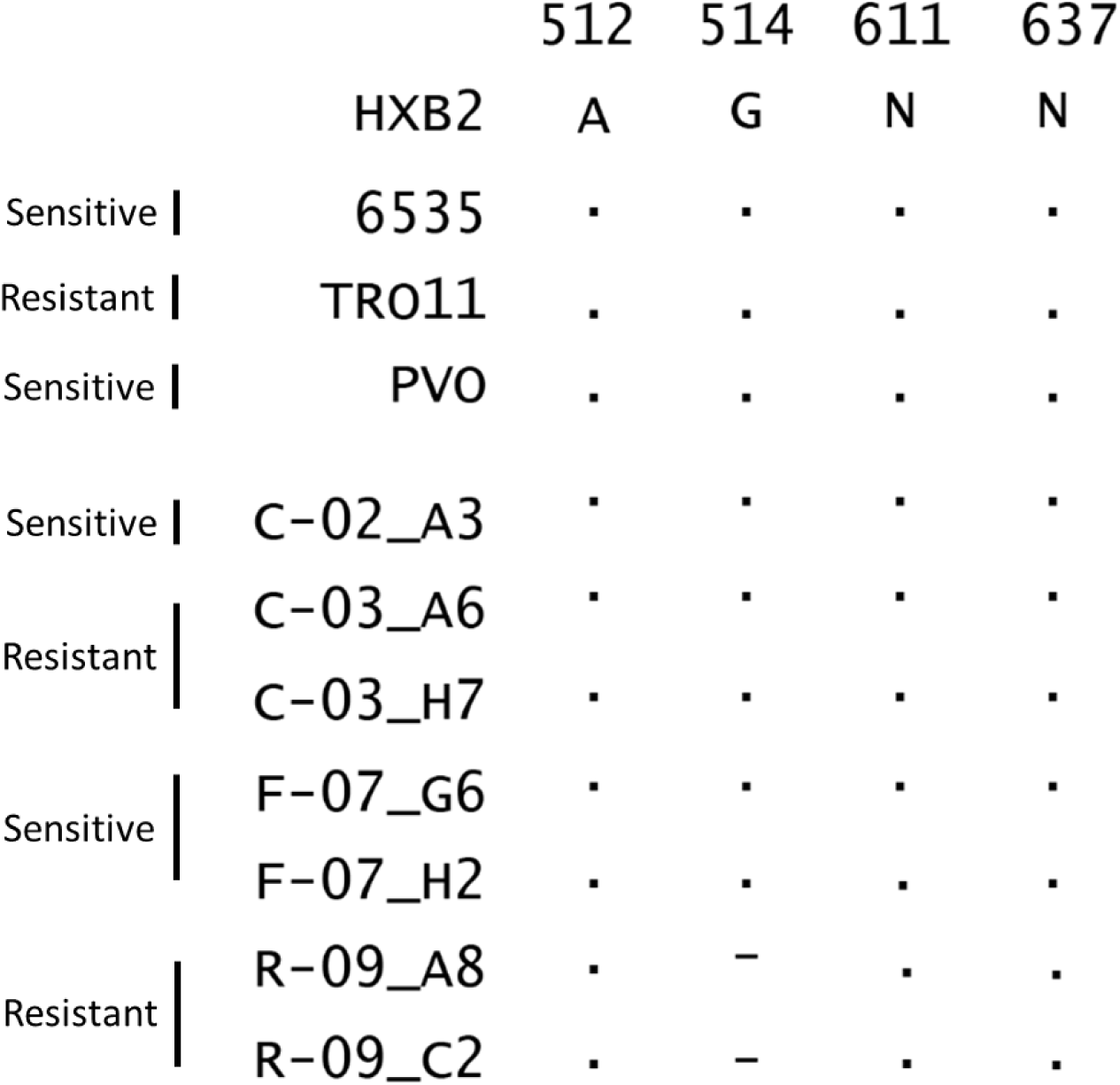
Missing Glycine Residue at HIV-1 Env position 514 renders R-09 sequences resistant to PGT-151. Key residues (A512, G514) and glycosylation sites (611, 637) involved in PGT-151 binding (Blattner, van Gils-16) are shown with donor Env residues and the control subtype B HIV-1 env sequences (6535, TRO11, PVO). Black dots represent homology to HIV-1 HXB2.

## Discussion

For HIV-1 Abs to be used therapeutically to clear low-level viremia and eliminate the infected cells producing viremia, it is important to ascertain the sensitivity of circulating virus to Ab binding and neutralization. We recently identified large clones of HIV-infected cells harboring infectious proviruses that were responsible for producing the low-level viremia observed (Halvas). This finding provided the unique opportunity to assess the sensitivity of viruses in plasma produced by infected cell clones to neutralization by autologous Ig or a broad panel of mAbs. From these four individuals on suppressive ART for a median of 14 years and exhibiting durations of detectable viremia (median of 19 c/mL) for a median of 4.85 years, seven unique PSVs were generated and tested (Fig 1) (Tables 1 and 2). The results of this work revealed complex patterns of sensitivity and resistance across and within donors, underscoring the diversity of virus and infected cells that persist even after long-term ART (Figs 2A-2D and S2-S7) (Table 3).

In contrast to the variable susceptibilities observed with our plasma-derived PSVs by monoclonal Abs (Table 3), contemporaneous autologous IgG from these donors showed little effect in neutralization to their respective PSVs with plasma-derived Envs (Figs 2A-2D), despite the presence of purified autologous Igs and specific Ig isotypes. The failure of autologous Igs to neutralize plasma virus may explain the non-suppressible viremia in these individuals, along with the persistence of clones with intact proviruses (Halvas). The presence of purified plasma-derived total Igs and specific Ig isotypes (i.e. IgG1-G4, IgA, and IgM) were confirmed by SDS-PAGE and quantitative ELISAs, respectively (S1 Fig) (S1 Table). A primary purified protein concentration of 500ug/mL was used as previously described (Esmaeilzadeh) and the inputted amount of IgG1-G4 into each neutralization assay was estimated to be about 40× higher than the primary concentration used for the panel of mAbs (S1 Table). The estimated amounts of IgA and IgM were found to be 6.2× and 12.4× higher, respectively than the primary concentrations used for the mAbs. The lack or absence of neutralization by autologous IgG could account for the episodes of low-level viremia previously observed in these participants (Halvas). The resistance of these viruses to neutralization by autologous Ig could is consistent with the inability to clear persistent viremia.

We also tested a panel of monoclonal Abs which target the major vulnerable sites in HIV-1 Env including the CD4bs, V3-glycan, V1V2, MPER, and the gp120-gp41 interface (Burton-16, Chen-16, Blattner, Huang-12, Xu, Kwon, Doria-Rose, Montero,). The patterns of sensitivity and resistance of individual HIV-Envs from the four donors were complex, with no one PSV being neutralization sensitive or less sensitive to all Abs tested (S2-S7 Figs) (Table 3). The R-09_C2 Env from was the least sensitive Env across all Abs tested (11 of 15 Abs) in contrast to F-07_H2 Env, the least resistant or most susceptible (two of 15 Abs). Furthermore, we observed no consistent pattern of resistance to neutralization with PSVs between donors and for the most part within donors, except for PGT151 (Table 3). In some cases, Envs for a specific donor were either all sensitive to a particular class of Abs (Donor F-07 for the V3-glycan Abs) or all less sensitive (Donor R-09 for the V1V2 Apex Abs) (Table 3). There were also instances in which the majority of PSVs tested were either less sensitive to a particular Ab (i.e. b12 and 2F5) or sensitive (i.e. N6/PGDM1400x10E8, 3BNC117, 10-1074, and 10E8) (S2-S7 Figs) (Table 3). Interestingly, the HIV-1 Envs produced by well-characterized infected cell clones (C-02_A3, C-03_H7, and R-09_A8) were found to still be sensitive to the majority (53%-73%) of Abs tested. As a result of this complexity, the unique neutralization profiles of these circulating viruses, even within a donor, will likely make it difficult to obtain therapeutic efficacy from just one Ab.

Previous studies have shown that longer variable loop lengths and/or PNGS are associated with resistance to Ab neutralization (van Gils-11, Doores, Goo). To assess the possible contributions of variable loop lengths and PNGS on neutralization efficiencies, analyses were performed to calculate i) the loop lengths, ii) the number of PNGS, and iii) the total glycans across Env. The mean variable loop lengths, the mean number of PNGS, and total glycans across the Envs from our donors were generally comparable to those in the controls (Table 4). A more detailed analysis of individual variable loops revealed some differences in the V1V2 and V5 regions as compared to the controls (Table 4). Of all the variable loops, the V1V2 regions from the donors displayed the most variability in lengths (range, 68-85 amino acids) and in the number PNGS (range, 7-9) (Table 4), but there was no obvious association of the pattern of either length or number of PNGS and neutralization. Interestingly, R-09_C2 Env had the shortest V1V2 loop length (68 amino acids) and the smallest number of PNGS (n = 7) but was highly less sensitive to neutralization by the Abs tested (Tables 3 and 4). Conversely, F-07_H2 Env had a V1V2 loop of intermediate length (76 amino acids) and seven PNGS was sensitive to neutralization. We also observed no significant association between i) the length of the V5 variable loop or ii) the number of V1 loop PNGS and neutralization. This was somewhat unexpected from what has been previously published (Ringe, van Gils-11, Doores, Goo), but the lack of association may be attributed to number of factors. These factors include: i) the engagement of other residues outside the variable loops, ii) the addition/subtraction of specific PNGS, and/or iii) the small sampling size (n = 7 Envs tested), limiting the power to detect associations.

Numerous mechanisms can contribute to resistance profiles associated with neutralization by anti-HIV-1 Abs. These can include specific amino acid substitutions in HIV-1 Env, which directly interact with the Ab and affect binding (Manhas, Zhou-19, Ringe, Caskey-15, Li-11) or more distal substitutions that can exert effects on binding and neutralization (Goo). Furthermore, addition, subtraction, or altered spatial location of a specific PNGS or changes in the overall electrostatic charge of specific HIV-1 Env domains can alter neutralization of HIV-1 Env by an Ab (Ringe, Wang). Complicating matters, evolution of resistance to neutralization by Abs can occur through numerous pathways generating complex patterns of substitutions (Otsuka). Therefore, despite the general lack of association between resistance to neutralization with either variable loop lengths, number of PNGS, and/or total number of glycans (Table 4), we opted to perform a deeper analysis of *env* sequences to determine the mechanism(s) associated with the resistance profiles we observed in donor-derived HIV-1 Envs.

We tested our PSVs against a number of Abs that targeted the CD4bs in HIV-1 Env of which all, except variant C-03_A6, were less sensitive (>2.5 µg/mL) to neutralization by b12 (S2 Fig) (Table 3). A number of amino acid substitutions were found in the *env* sequences, which have been implicated in resistance to b12 neutralization (S2 Table) (Wu-09), but these substitutions did not explain the discordant neutralization sensitivities observed across C-03 donor HIV-Envs with identical amino acids at these positions. This finding suggested that other residues may account for the variation observed with b12 neutralization sensitivities between C-03_A6 and C-03_H7. We observed a total of 21 amino acid differences which included i) loss or addition of a single amino acid, ii) loss of PNGS in C-03_H7, iii) creation of new PNGS, and iv) drastic changes in side chain composition (i.e. nonpolar or polar versus positively charged, hydrophobic versus positively charged, or aromatic ring versus hydrophobic) (S3 Table). It is plausible that some of these mutations affect quaternary structure, leading to poor binding of b12 (Wu-09).

Many VRC01 resistance mutations have been previously identified (Zhou-19, Otsuka). Out of the five individuals studied here only one donor, R-09, harbored Env sequences that were less sensitive to the CD4bs bNAb VRC01 (S2 Fig) (Table 3). Both R-09 Env sequences (R-09_A8 and R-09_C2) had an additional PNGS at N128, an extended V5 region with an additional glycan, and the introduction of a neutral Q residue at position 476, which typically is occupied by positively charged K/R (Fig 3). Given the known role of PNGS to mask epitopes, the addition of a PNGS at residue 128 could contribute to the VRC01-resistance observed with donor R-09 HIV-1 Envs. Furthermore, residue 476 at the base of V5 region, which is a contact for VRC01, is a charge neutral glutamine, differing from the more dominant positively charged lysine or arginine at this position (Fig 3). Changes that affect charges are interesting, given the potential for attraction/repulsion of nearby regions on gp120, which could affect epitope exposure and/or Ab binding.

Other mutations outside of the VRC01 epitope are also known to influence VRC01 resistance. For example, disruption of the gp120 glycan N262, which is outside of the CD4bs and not a VRC01 contact residue, decreased VRC01 binding to the subtype B Env JRCSF to <33% of wild type (Huang-16). While disruption of N262 was not observed in our donors, we did find that both R-09 HIV-1 Envs harbored at least one mutation in gp41 (K683R) that was previously identified as affecting CD4bs Ab susceptibility (Fig 5) (Bricault).

The VRC01-resistant Envs, R-09_A8 and R-09_C2, were also less sensitive to the V1V2 apex bNAbs PG9, PG16 and PGT-145 (Fig S4) and possibly explained by the Q residue at position 160 disrupting a common PNGS that is required for PG9 and PG-9 related Ab binding (Fig 4B). Interestingly, in donor R-09 HIV-1 Envs, we observed a PNGS at position 128 possibly contributing to VRC01-resistance, and also possibly implicated with resistance to PG9 (Fig 3) (Doria-Rose). Given that the R-09 Envs were the most resistant to the Abs tested in this study, these Envs likely have substitutions that disrupt Ab contact sites and/or may ‘mask’ neutralizing epitopes. The conformational flexibility of the HIV-1 Env trimer, in both the CD4 bound and unbound states can affect Ab binding (Korkut). Therefore, changes in Env such as extended variable loops or the number of PNGS can likely impact Ab binding due to physical changes associated with Env epitopes.

It was recently found that a high number of PNGS in V1 are associated with 10E8, 4E10 and 2F5 neutralization sensitivity (Bricault). We did not observe any of these associations in our data set (Data not shown) and neutralization sensitivities were variable among and within donors (S5 Fig) (Table 3). The MPER epitope signature K683R is also associated with resistance to CD4bs and V2 bNAbs (Bricault). Interestingly, variants R-09_A8 and R-09_C2, which are less sensitive to the CD4bs Ab VRC01 (S2 Fig) (Table 3) and V1V2 Abs PG9, PG16 and PGT-145 (S4 Fig) (Table 3) have an Arg at position 683 (Figs 5) (Table 3). This Gp41 mutation could be a mechanism involved in modifying CD4bs and V1V2 neutralization of donor R-09 Envs. Finally, R-09_C2 lack of sensitivity to the tri-specific Ab could be attributed to the aforementioned changes to the CD4bs, V1V2 apex, and gp41 regions of Env, but do not explain R-09_A8 Env sensitivity and suggest that other residues are also involved (S7 Fig) (Table 3).

Variants C-03_H7 and C-03_A6 lacked sensitivity to PGT-121 neutralization, whereas the remaining Envs tested, were susceptible to PGT-121 (S3 Fig) (Table 3). It has been well established that the net charge of the V3 region as well as charge of particular residues are important for 3D orientation of V3 on Env trimer and co-receptor binding (Chandramouli). Variants C-03_H7 and C-03_A6 had a neutral glutamine residue at position 330, while most of the PGT-121 sensitive Env sequences had a positively charged histidine at this position (Fig 4A). Given the importance of V3 charge, this change could be altering the 3D orientation of V3, making it more difficult for PGT-121 to neutralize. We observed no distinct amino acid changes in V3 for R-09_C2 that would account for PGT-128 reduced sensitivity to neutralization and suggests that residues outside V3 were likely contributing to resistance.

An effective HIV-1 cure will only be achieved when circulating and latent HIV-1 are eliminated or prevented from reseeding a spreading infection. A popular approach to achieve this has been the use of mAbs that target circulating Env as well as Env produced by reactivated latently infected cells. The diversity of Env sensitivity to neutralization by mAbs, that we have shown in a limited number of donors, indicates in-depth analyses of HIV-1 Env sequences are needed to identify the combinations of mAbs that are most effective in neutralizing and clearing virus (Yu, 2019). Combinations of several antibodies or multi-specific single antibodies hold promise toward this goal. Alternatively, *in vivo* expression of a very broadly neutralizing and long-lasting inhibitor targeting viral entry could potentially prevent viral rebound after ART is stopped. In this regard, eCD4-Ig displayed low neutralization IC50 values and Ab-dependent cell-mediated cytotoxicity and was expressed in vivo sufficiently to prevent infection in non-human primate models (Gardner). Whether all infected cells harboring replication competent proviruses can be eliminated remains to be determined.

## Method details

### Study participants

Plasma samples were obtained, following written informed consent, from donors with non-suppressible viremia while on ART under an appropriate human research protocol (PRO10070203) approved by the University of Pittsburgh’s institutional review board. Non-suppressible viremia was defined as plasma HIV-1 RNA >40 copies/ml for more than six months after long-term suppression on ART. Single genome sequencing (SGS) of HIV-1 *env* was performed on four PWL (C-02, C-03, F-07, and R-09) (Table 1) (Palmer).

### HIV-1 plasma RNA quantification

Plasma HIV-1 RNA copies/mL was measured by the COBAS Ampliprep/COBAS TaqMan, v2.0 assay (TMv2.0) (Roche, Basel, CH) or M2000 RealTime HIV-1 Viral Load Assay (Abbott Molecular, Des Plaines, IL, USA). Alternatively, plasma HIV-1 RNA was quantified in batch on frozen plasma processed from peripheral blood using a single copy assay as was previously described (Cillo). In brief, EDTA anticoagulated plasma was ultra-centrifuged at 170,000 g for 30 min, HIV-1 RNA was extracted at 42 C^0^ for 1h using 3M guanidinium hydrochloride, followed by 6M guanidinium thiocyanate and 600 g/mL of glycogen (Roche, Indianapolis, IN) at 42^0^C for an additional 10 min. HIV-1 RNA was precipitated in 100% isopropanol by centrifugation at 21,000g for 10 min, the RNA pellet washed in 70% ethanol, air dried, and resuspended in 5mM Tris-HCl with 1M dithiothreitol and 1,000 U/mL of RNasin (Promega, Maddison, WI). A replication-competent avian leukosis virus (ALV) long terminal repeat with a splice adaptor (RCAS) was spiked into each sample and used as a quantity of internal standard.

HIV-1 RNA and RCAS was quantified using a two-step qRT-PCR by reverse transcribing cDNA that was then inputted as template into a real-time PCR using the LightCycler 480 Probes Master ready-made master mix with the addition of primers and probes targeting HIV-1 RNA and RCAS. Copy numbers per mL were quantified against a standard curve generated by highly characterized HIV-1 transcripts, which were serial diluted 3-fold to 1 copy per qRT-PCR. A number of controls were run in parallel with the samples tested including i) a negative human plasma sample, ii) low-copy-number plasma with 5 HIV-1 RNA copies/mL (Rush University, Virology Quality Assurance Lab, Chicago, IL, USA), and iii) a no reverse transcriptase control to exclude the presence of HIV-1 DNA (Cillo).

### Cells, media, and plasmids

Human embryonic kidney (HEK)-293T cells (NIH AIDS Reagent Program (NIH ARP), Manassas, VA, USA) were used to generate pseudovirus (PSV) and TZM-bl cells (National Institute of Health AIDS Reagent Program (NIH ARP), Bethesda, MD, USA) were used as target cells in neutralization assays. Both cell lines were cultured in Dulbecco’s Modified Eagle Medium (DMEM; Gibco, Amarillo, TX, USA) supplemented with 10% fetal bovine serum, antibiotics (60 units/ml penicillin and 60 µg/ml streptomycin) and 1x GlutaMax (Gibco, Waltham, MA, USA). The following subtype B *env* expression plasmids were obtained from the NIH ARP: 6535 (tier 1B), TRO11 (tier 2), and PVO (tier 3) that have above average, moderate and low sensitivity to Ab neutralization respectively (Seaman).

### Plasma viral RNA extractions and cDNA synthesis

Plasma RNA extractions and subsequent cDNA synthesis were performed as reported (Palmer, Halvas) with minor modifications. In brief, virions were pelleted from plasma at 24 000 x g for 1h at 4°C, lysed in buffer [final concentrations; 3.6M guanidine thiocyanate, 0.13M DTT, 0.67 mg/ml glycogen, 34 mM N-Lauroylsarcosine, 34 mM sodium citrate (Sigma-Aldrich, St. Louis, MO, USA)], precipitated with 100% isopropanol (Sigma-Aldrich, St. Louis, MO, USA), and washed with 70% ethanol. The final pellets were re-suspended with 20 µl of cold 5 mM Tris-HCl pH 8.0 (Sigma-Aldrich, St. Louis, MO, USA). The RNA was then denatured at 65°C for 10 min with 0.5 mM dNTPs and the *env* PCR1 reverse primer ENV_R1 (5’-TTGCTACTTGTGATTGCTCCATGT-3’, HXB2 positions 8469-8492) (GENEWIZ, South Plainfield, NJ, USA) at 0.25 µM. The entire RNA sample was used immediately for reverse transcriptase PCR (RT-PCR) to produce cDNA with Superscript III (Invitrogen, Carlsbad, CA, USA) in a total reaction volume of 50 µl [final concentrations;1X RT buffer, 5 mM DTT, 5 mM MgCl2, 0.4 U/uL RNaseOut, 10 U/uL Superscript III]. The RT-PCR reaction was incubated as follows: 50°C for 1 h, 55°C for 1 h, 70°C for 15 min, and stored at -20°C until downstream use.

### Envelope single genome amplification

HIV-1 *env* cDNA was serially diluted (1:2 or 1:4) to an endpoint in 5mM Tris-HCl, pH 8.0 (Sigma-Aldrich, St. Louis, MO, USA) and distributed in 96-well semi-skirted PCR plate (USA Scientific, Ocala, FL, USA) such that PCR positive reactions occurred in <30% of wells, which by Poisson distribution is estimated to result in one cDNA template per positive reaction (Palmer). No template controls were included on each reaction plate. Nested PCR was used to amplify HIV-1 *env* in a 10 µl reaction [Final concentrations;1X High Fidelity PCR buffer, 2 mM MgSO4, 0.2 mM dNTPs, 0.2 µM each primer, 0.04 U/µl Platinum Taq High Fidelity polymerase (Invitrogen, Carlsbad, CA, USA)]. The following outer primers were used for the first round of PCR: ENV_F1 (5’-TAGAGCCCTGGAAGCATCCAGGAAG-3’, HXB2 positions 5409-5433) and ENV_R1 (5’-TTGCTACTTGTGATTGCTCCATGT-3’, HXB2 positions 8469-8492) with the following touchdown PCR cycling parameters (PCR1: 94°C, 2 min; 5 cycles at 94°C, 30s; 64°C, 30s; 68°C, 4 min followed by 5 cycles at 94°C, 30s; 62°C, 30s; 68°C, 4 min followed by 5 cycles at 94°C, 30s; 60°C, 30s; 68°C, 4 min followed by 5 cycles at 94°C, 30s; 58°C, 30s; 68°C, 4 min followed by 15 cycles at 94°C, 30s; 55°C, 30s; 68°C, 4 min; and a final extension at 68°C, 10 min). PCR1 reactions were diluted 1:9 in 5mM Tris-HCl, pH 8.0 (Sigma, St. Louis, MO, USA) and 2 µl were transferred to PCR2 plates. The following inner primers were used for the nested PCR: ENV_F2 (5’-CACCTTAGGCATCTCCTATGGCAGGAAGAAG-3’, HXB2 positions 5509-5539) and ENV_R2 (5’-GTCTCGAGATACTGCTCCCACCC-3’, HXB2 positions 8438-8460) with the following PCR2 touchdown cycling conditions: (94°C, 2 min; 5 cycles at 94°C, 30s; 69°C, 30s; 68°C, 4 min followed by 5 cycles at 94°C, 30s; 67°C, 30s; 68°C, 4 min followed by 5 cycles at 94°C, 30s; 65°C, 30s; 68°C, 4 min followed by 5 cycles at 94°C, 30s; 63°C, 30s; 68°C, 4 min followed by 15 cycles at 94°C, 30s; 59.5°C, 30s; 68°C, 4 min; and a final extension at 68°C, 10 min). The nested HIV-1 *env* PCR amplicon (∼3 kb) was purified using solid phase immobilization beads (SPRI; KAPA Biosystems, Wilmington, MA, USA), diluted in 40uL 5mM Tris HCl, pH 8.0 (Sigma-Aldrich, St. Louis, Mo, USA), and stored at -20^0^C until sequencing.

### Sequencing, assembly, and phylogenetic analysis

HIV-1 *env* PCR amplicons were sequenced using the Illumina MiSeq platform per the manufacturer’s recommendations (Illumina, San Diego, CA, USA). Briefly, ∼50ng of each PCR product was enzymatically fragmented with Fragmentase (New England Biolabs (NEB), Ipswich, MA, USA), the fragmented amplicons were end-repaired, A-tailed, and the adapter ligated to generate Illumina libraries of 300-600 bp. The libraries were further amplified for 12 cycles and at each step of library construction, samples were purified by size-selection with KAPA Pure Beads (KAPA Biosystems, Wilmington, MA, USA). The final library sizes were validated with the Perkin Elmer GX Touch 24 LabChip bioanalyzer using the 3K DNA assay and chip. Sequencing libraries were run using a 500 cycle Illumina MiSeq Nano v2 kit and flow cell. Reference-guided alignments of sequencing reads were performed using BWA-MEM (Li H 2013) on Sequencher 5.4 (Gene Codes, Ann Arbor, USA). HIV-1 *env* sequences were aligned using Clustal and Molecular Evolutionary Genetics Analysis (MEGA X) was used to generate neighbor-joining phylogenetic trees (Thompson, Kumar).

### HIV-1 *env* cloning

HIV-1 *env* PCR products were gel extracted and ligated into the pCDNA3.1 directional TOPO vector (Invitrogen, Carlsbad, CA, USA). Top10 bacterial cells were transformed with the ligation reactions and colonies were used for plasmid minipreps then screened by *Xho I* and *Hind III* digestions for the presence of the insert. Plasmid preps containing an insert band of the expected size were confirmed by Sanger sequencing. Any point mutations in plasmid sequences were corrected with the QuikChange II XL Site-Directed Mutagenesis kit (Agilent, Santa Clara, CA, USA) to match the MiSeq *env* PCR sequence.

### Pseudovirus production

Pseudoviruses (PSVs) were generated by transient co-transfection of 293T cells with an *env* expressing plasmid and the HIV-1 luciferase reporter plasmid pNL4-3.Luc.R-.E- (NIH ARP) at a 1:2 ratio using polyethylenimine (25 kDa; Polysciences, Warrington, PA, USA) as the transfection reagent as reported (Pantophlet, Manhas). Supernatant containing pseudovirus was harvested 3 days post transfection, aliquoted and stored at -80°C.

### Autologous plasma antibody isolation

Autologous IgG was isolated in 3 phases involving i) heat inactivation of samples, ii), bead preparation, and iii) autologous IgG purification. Each serum sample was initially decomplemented by incubation at 56^0^C for 30 minutes followed by centrifugation at 10K rpm for 20 minutes (2 per participant). Protein A beads were washed per manufacturers recommendations (Bio-Rad, Hercules, CA, USA). Samples were prepared for purification by adding 450uL of washed beads to each tube containing 440uL of decomplemented serum to a total volume of 890uL. 890uL of 1X PBS was added to 890uL of sample, which was incubated on a spinning wheel overnight at 4^0^C.

The following day, the serum-bead mixture was transferred to a Bio-Rad column described above and washed with >4mL of 1X PBS by gravity flow using 1mL of 1X PBS and resuspension of beads each time, avoiding bead loss and drying. To elute IgG from beads, 1X PBS was removed from beads using gravity flow avoiding drying of beads, column was capped, and beads were resuspended in 450uL of Elution Buffer, incubated mixture for 1 minute at ambient temperature, and eluted and captured bound IgG by gravity flow in a new tube with neutralizing Tris-Base pH8. Elution steps were repeated 4 more times. If the concentration of IgG was detectable in 5^th^ elution, the elution process was repeated.

Eluted IgG underwent buffer exchange to 1X PBS using a 30kDa Filter Unit. All elution fractions from the same serum sample were pooled, added to a pre-washed 30kDa Filter Unit filled with 1X PBS, centrifuged for 20 minutes at 3.5K rpm, leaving a remaining volume of <1mL. and resuspended the small IgG volume by mixing using a pipettor, avoiding bubbles, and flushing membrane with the IgG volume. The filter was refilled with additional 1X PBS and steps repeated three more times. During the final buffer exchange, the final volume in 1X PBS remaining was <400uL. Membranes were flushed, remaining volume transferred to a 1.5mL filter tube (0.22um), volume adjusted to 400uL with 1X PBS, and filter centrifuged 4K rpm for 10 minutes, so that the entire volume passed through the filter. The filter was removed and concentration measured using De Novix DS-11 spectrophotometer (De Novix, Wilmington, DE, USA) at 280nm. Isolated IgG was transferred to new sterile and cryogenic screw cap tube and stored at - 80^0^C until used in neutralization experiments.

### Detection of total immunoglobulins (Ig) by sodium dodecyl sulfate polyacrylamide gel electrophoresis

Sodium dodecyl sulfate polyacrylamide gel electrophoresis (SDS-PAGE) and enzyme-linked immunosorbent assay (ELISA) was used to confirm the presence of total immunoglobulins (Ig) and the IgA, IgG, and IgM isotypes, respectively. 3ug of purified protein sample in SDS-PAGE loading buffer (Invitrogen, Waltham, MA, USA) in the presence or absence of 50mM DTT (Invitrogen, Waltham, MA, USA) were boiled for 10 minutes, resolved at 135V for 35 minutes on a NuPAGE 10% Bis-Tris gel (Invitrogen, Waltham, MA, USA) and stained using Coomassie Blue (Invitrogen, Waltham, MA, USA).

### Quantitative enzyme-linked immunosorbent assay of IgG (1-4), IgA, and IgM

The human IgG, IgA, and IgM ELISA kits (Thermo Fisher, Waltham, MA, USA) were used as recommended by the manufacturer to quantify IgG1, IgG2, IgG3, and IgG4 (IgG1-G4), IgA, and IgM isotype concentrations in plasma and purified protein from plasma. Briefly, purified plasma was diluted to 0.375 ug/mL, 2.75 ug/mL, and 3.6 ug/mL separately for IgG, IgA, and IgM tests. The pre-coated microwell strips were washed twice with washing buffer at 400 uL/well. The standard IgG, IgA, and IgM were diluted on the microwell plate with 2-fold serial dilution from 100 ng/mL for IgG and IgA, and 1000 ng/mL for IgM. 100 uL/well of the diluted sample was added to the sample wells. 50 uL of diluted HRP-conjugated anti-IgG, anti-IgA, and anti-IgM antibodies were separately added to all wells, then the microwell plate was incubated at room temperature for 1 hour with shaking. After incubation, the microwell strips were washed four times with washing buffer. The enzyme reaction was developed by adding 100 uL of TMB substrate solution and incubating at room temperature for 30 minutes to develop color, then stopped by stop solution. The absorbance of each microwell was recorded at 450 nm. Experiments were performed in duplicate. To determine the concentration of Ig subtypes in plasma and purified plasma, the average absorbance values of each set of duplicate standards and samples were calculated. The standard curve was created by plotting the mean absorbance for each standard concentration using a binomial equation. The sample concentrations were calculated from the standard curve by multiplying with the dilution factor.

### Monoclonal antibodies

The following monoclonal antibodies (mAbs) were tested: directed to the i) CD4 binding site [CD4bs; b12 (Barbas), VRC01 (Wu), 3BNC117 (Klein), and LSEVh-LS (Chen)]; ii) V1V2 apex [PG9 (Walker-09), PG16 (Walker-09), and PGT-145 (Walker-11)]; iii) V3-glycan [PGT-121 (Walker-11), PGT-128 (Walker-11), and 10-1074 (Mouquet)]; iv) gp120-gp41 interface [PGT151 (Blattner)]; v) the membrane-proximal external region (MPER) of gp41 [2F5 (Buchacher), 4E10 (Buchacher), and 10E8 (Huang-12)]; and vi) a tri-specific antibody [N6/PGDM1400x10E8 (Xu)] directed to the CD4bs, V1V2 apex, and MPER binding sites.

### Virus neutralization assay

Neutralization assays were performed as reported (Manhas, Richman). Briefly, pseudoviruses were incubated with antibody dilutions for 1 h at 37°C, then PSV/Ab mixtures were added to TZM-bl target cells in the presence of Diethylaminoethyl-Dextran (DEAE-dextran; Sigma-Aldrich, St. Louis, MO, USA). Target cells were cultured for 3 days after which luciferase activity was measured on a Luminoskan Accent (Thermo Fisher Scientific, Waltham, MA, USA) or Victor Nivo (Revvity, Waltham, MA, USA) using the Britelite Plus Reporter Gene Assay System (Perkin Elmer, Waltham, MA, USA). Percentage neutralization was calculated relative to cell-only and virus-only controls. Neutralization curves were fitted using GraphPad PRISM v10.0.0 (GraphPad Software, LLC, San Diego, CA, USA).

Neutralization assays to test autologous IgG were performed as reported above for the monoclonal antibodies (Manhas, Richman) with the following modifications. Participant-derived PSVs were tested against their respective contemporaneous autologous IgGs starting with a primary autologous IgG concentration of 500ug/mL. Autologous IgG was serially diluted 1:3 (range of autologous IgG concentrations; 500 to 0.229ug/mL). Neutralization assays controls tested in parallel included testing autologous IgG against PSVs 6535, TRO.11, and PVO as well as a virus with no autologous IgG. Assay run controls included testing PSVs TRO.11 and PVO against the monoclonal antibody 10-1074. Monoclonal antibody 10-1074 was serially diluted 1:5 (range of 10-1074 concentrations; 20 to 0.009ug/mL). Neutralization curves were generated by GraphPad PRISM v10.0.0 (GraphPad Software LLC, San Diego, CA, USA).

## Data availability

Sequences have been submitted to the GenBank database (accession numbers OR791051-OR791082).

## ACKNOWLEDGEMENTS

We thank the donors for their participation in this study. We thank the Dimitrov lab at the University of Pittsburgh for LSEVh-LS as well as the NIH AIDS Reagent Program (NARP) for all other Abs used in this study. We also thank the NARP for the 6535, TRO11 and PVO *env* expression plasmids used in this study. We are also grateful to Gilead for providing some of the Abs used in this study and the Virology and Persistence Core of the Rustbelt Center for AIDS Research (P30 AI036219) for performing the neutralization assays utilizing the participant-derived autologous IgGs. We thank Lorraine Pollini for proofreading, formatting, and submission of this manuscript.

## Author Contributions

**Conceptualization:** John W. Mellors.

**Data curation:** Savrina Manhas, Elias K. Halvas.

**Formal analysis:** Savrina Manhas, Kerri J. Penrose, Xiaojie Chu, Wei Li, John W. Mellors, Elias K. Halvas.

**Funding acquisition:** Elias K. Halvas, John W. Mellors, Mary F. Kearney.

**Investigation:** Savrina Manhas, Joseph P. Brooker, Cory Shetler, Kerri J. Penrose, Divya S. Jaiswal, Xiaojie Chu, Wei Li, Elias K. Halvas.

**Methodology:** Savrina Manhas, Joseph P. Brooker, Cory Shetler, Kerri J. Penrose, Divya S. Jaiswal, Xiaojie Chu, Wei Li.

**Project administration:** Elias K. Halvas, John W. Mellors, Mary F. Kearney.

**Resources:** Elias K. Halvas, John W. Mellors, Mary F. Kearney.

**Software:** Savrina Manhas, Kerri J. Penrose, Wei Li, Mary F. Kearney, John W. Mellors, Elias K. Halvas.

**Supervision:** Elias K. Halvas, John W. Mellors.

**Validation:** Elias K. Halvas, John W. Mellors, Mary F. Kearney.

**Visualization:** Divya S. Jaiswal, Xiaojie Chu, Wei Li.

**Writing – original draft:** Savrina Manhas, Elias K. Halvas, John W. Mellors

**Writing – review & editing:** Savrina Manhas, Joseph P. Brooker, Cory Shetler, Kerri J. Penrose, Divya S. Jaiswal, Xiaojie Chu, Wei Li, Mary F. Kearney, John W. Mellors, Elias K. Halvas.

