## Supplemental Figures and Tables for "HIV-1 envelopes from virions that persist in plasma on antiretroviral therapy show reduced susceptibility to autologous immunoglobulins and variable sensitivity to broadly neutralizing monoclonal antibodies"

Variable autologous and monoclonal antibody sensitivity profiles in people with HIV

**Savrina Manhas[1], Joseph P. Brooker[1], Cory Shetler[1], Kerri J. Penrose[1], Divya S. Jaiswal[2], Xiaojie Chu[2], Wei Li[2], Mary F. Kearney[3], John W. Mellors[1], Elias K. Halvas[1*]**

**1** Division of Infectious Diseases, Department of Medicine, University of Pittsburgh School of Medicine, Pittsburgh, Pennsylvania, 15261, United States of America

**2** Center for Antibody Therapeutics, Division of Infectious Diseases, Department of Medicine, University of Pittsburgh School of Medicine, Pittsburgh, Pennsylvania, 15261, United States of America

**3** HIV Dynamics and Replication Program, National Cancer Institute, Frederick, MD 21702, United States of America

Present address: Elias K. Halvas, Division of Infectious Diseases, School of Medicine, University of Pittsburgh, Scaife Hall, Suite 807A, 3550 Terrace Street, Pittsburgh, PA 15261

**Supporting information**

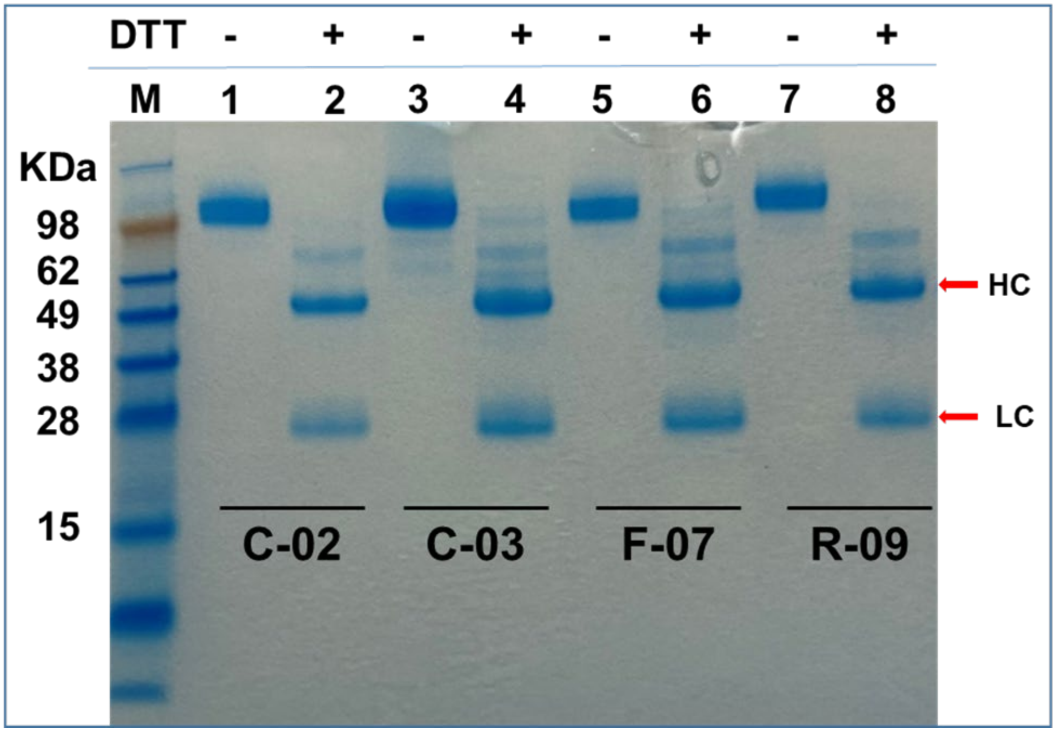

**S1 Fig. Sodium dodecyl sulfate-polyacrylamide gel electrophoresis (SDS-PAGE) of purified plasma immunoglobulins (Igs).** Autologous Igs from donors C-02, C-03, F-07, and R-09 were purified from plasma. 5-10 ug of purified proteins was resolved in SDS-PAGE loading buffer in both the presence and the absence of reducing reagents dithiothreitol (50 mM). Samples were boiled for 10 mins before loading onto SDS-PAGE. Gel was stained by Coomassie Blue.

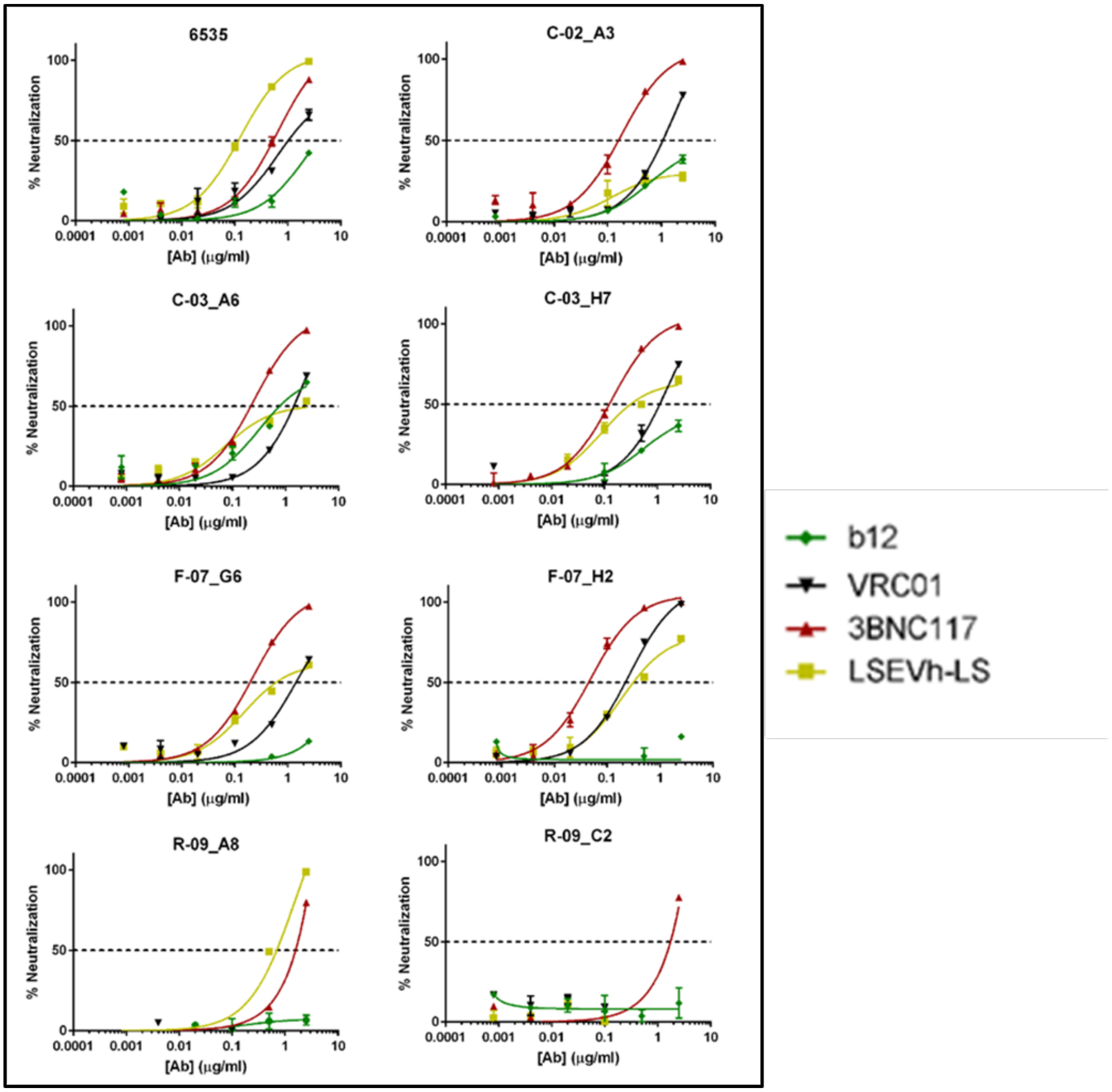

**S2 Fig. Neutralization sensitivities for CD4 binding site (CD4bs) monoclonal antibodies (mAbs) LSEVh-LS, 3BNC117, VRC01, and b12.** Donor C-02, C-03, F-07, and R-09 HIV-1 envelopes were expressed as pseudovirions and incubated with CD4bs mAbs (VRC01, 3BNC117, LSEVh-LS, or b12) for 1 h then added to TZM-bl target cells. Luciferase was measured after 3 days. Percentage neutralization was calculated relative to cell-only and virus-only controls. The tier 1b subtype B pseudovirus 6535 is used as a neutralization control. Neutralization curves fitted by GraphPad PRISM v10.0.0.

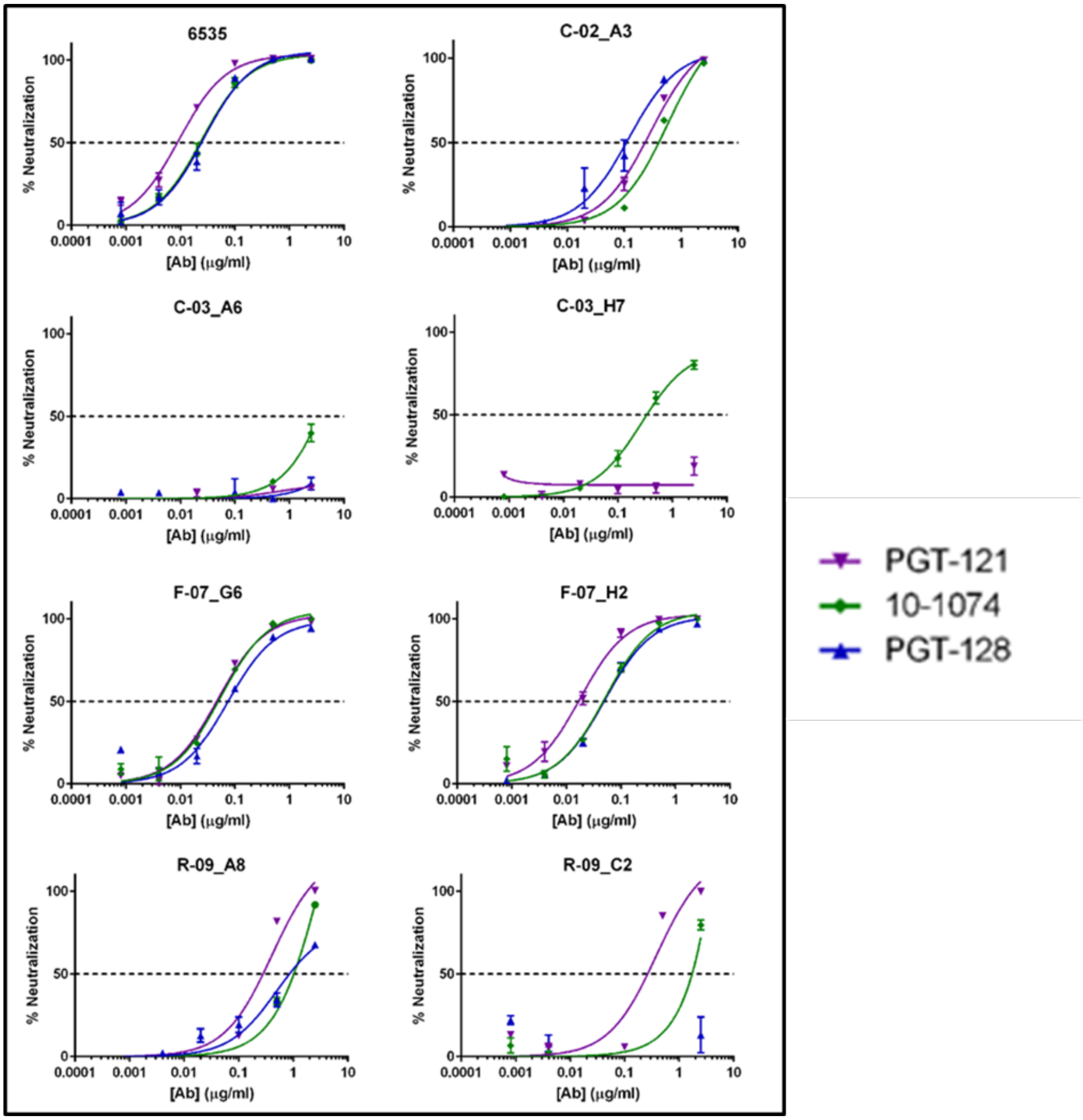

**S3 Fig. Neutralization sensitivities for V3-glycan monoclonal antibodies (mAbs) PGT-128, PGT-121, and 10-1074.** Donor C-02, C-03, F-07, and R-09 HIV-1 envelopes were expressed as pseudovirions and incubated with V3-glycan mAbs (PGT-128, PGT-121, or 10-1074) for 1 h then added to TZM-bl target cells. Luciferase was measured after 3 days. Percentage neutralization was calculated relative to cell-only and virus-only controls. The tier 1b subtype B pseudovirus 6535 is used as a neutralization control. Neutralization curves fitted by GraphPad PRISM v10.0.0.

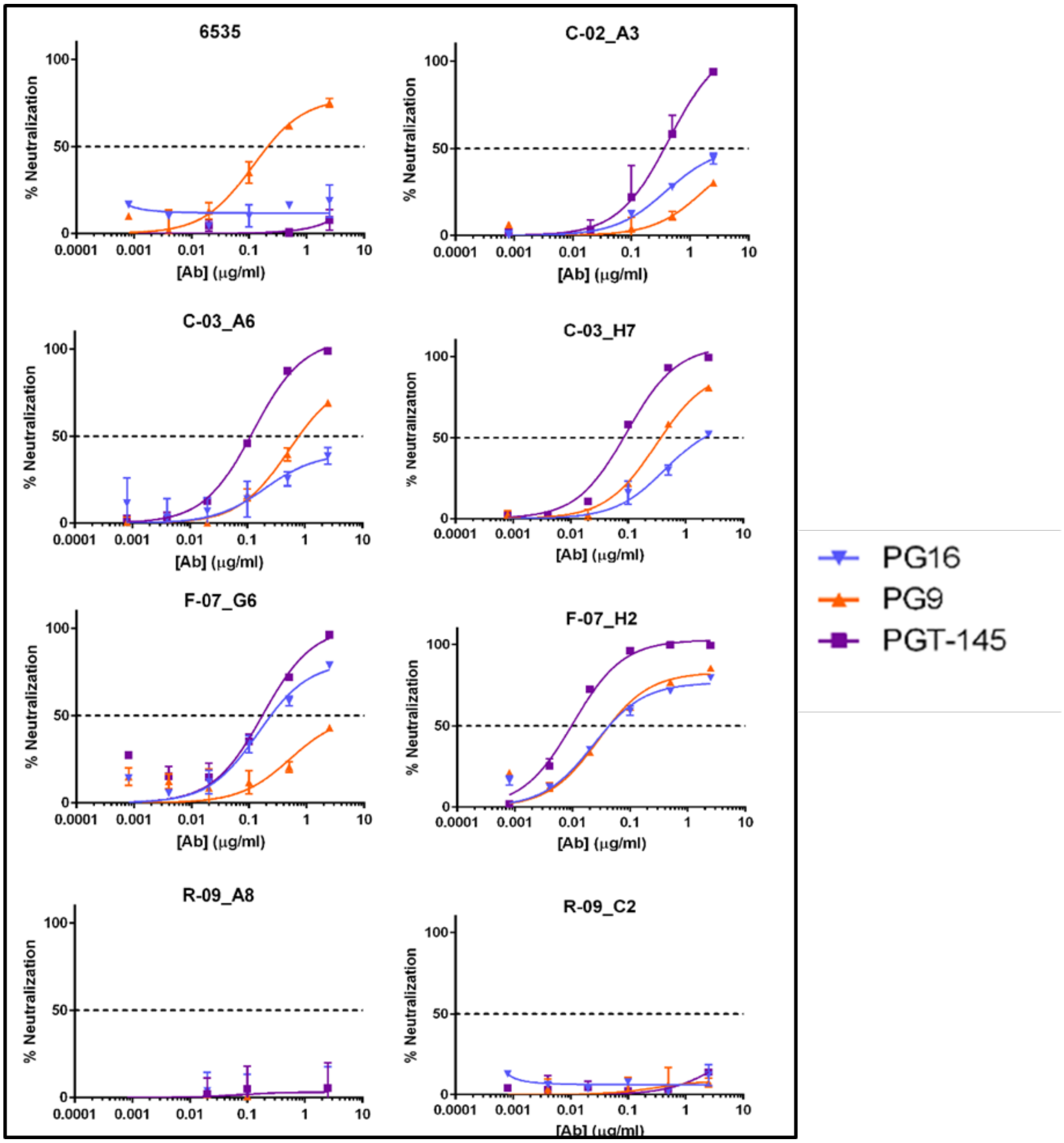

**S4 Fig. Neutralization sensitivities for Apex V1V2 monoclonal antibodies (mAbs) PG16, PG9, and PGT-145.** Donor C-02, C-03, F-07, and R-09 HIV-1 envelopes were expressed as pseudovirions and incubated with Apex V1V2 mAbs (PG16, PG9, or PGT-145) for 1 h then added to TZM-bl target cells. Luciferase was measured after 3 days. Percentage neutralization was calculated relative to cell-only and virus-only controls. The tier 1b subtype B pseudovirus 6535 is used as a neutralization control. Neutralization curves fitted by GraphPad PRISM v10.0.0.

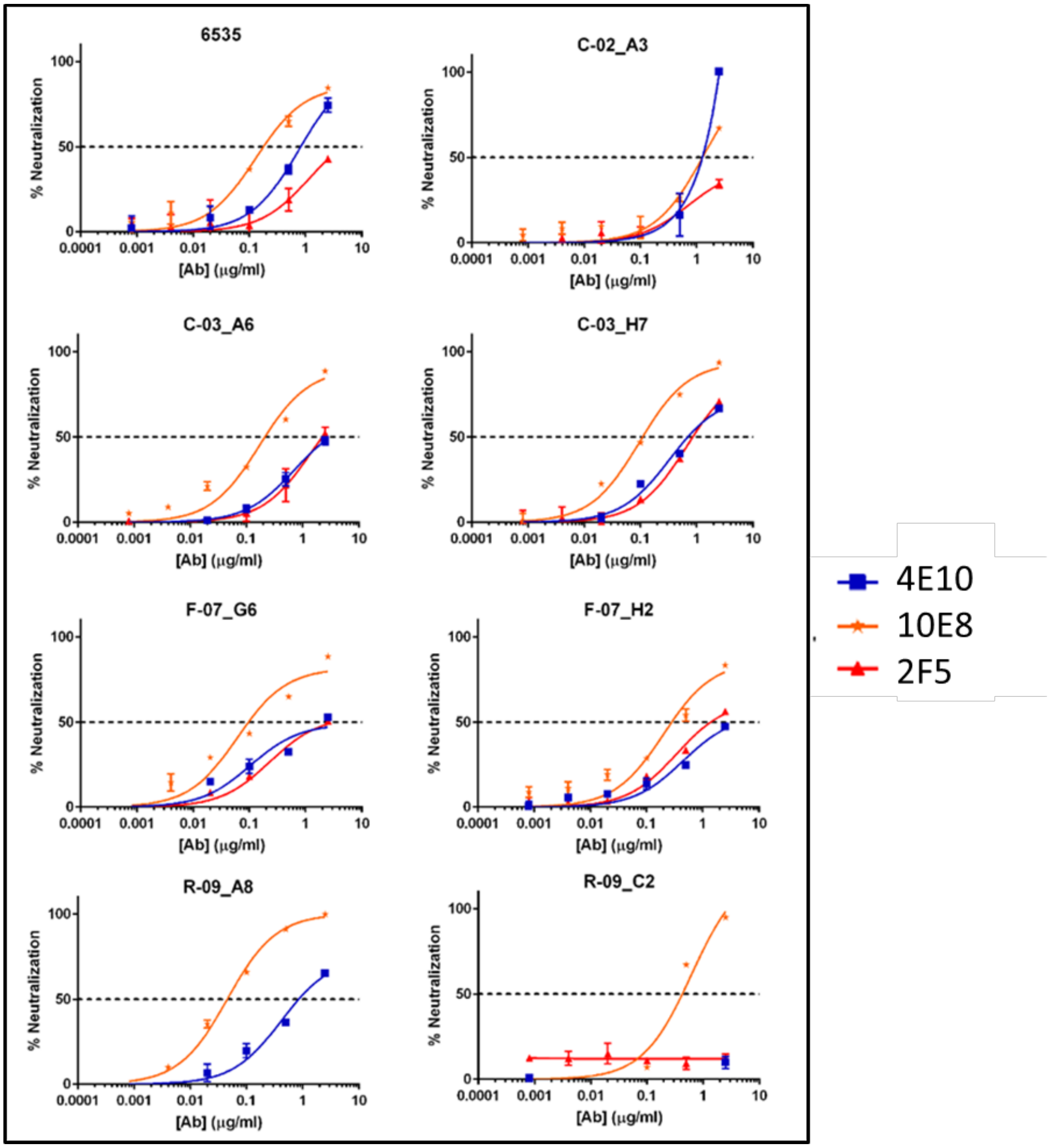

**S5 Fig. Neutralization sensitivities for gp41 monoclonal antibodies (mAbs) 4E10, 10E8, and 2F5.** Donor C-02, C-03, F-07, and R-09 HIV-1 envelopes were expressed as pseudovirions and incubated with gp41mAbs (4E10, 10E8, or 2F5) for 1 h then added to TZM-bl target cells. Luciferase was measured after 3 days. Percentage neutralization was calculated relative to cell-only and virus-only controls. The tier 1b subtype B pseudovirus 6535 is used as a neutralization control. Neutralization curves fitted by GraphPad PRISM v10.0.0.

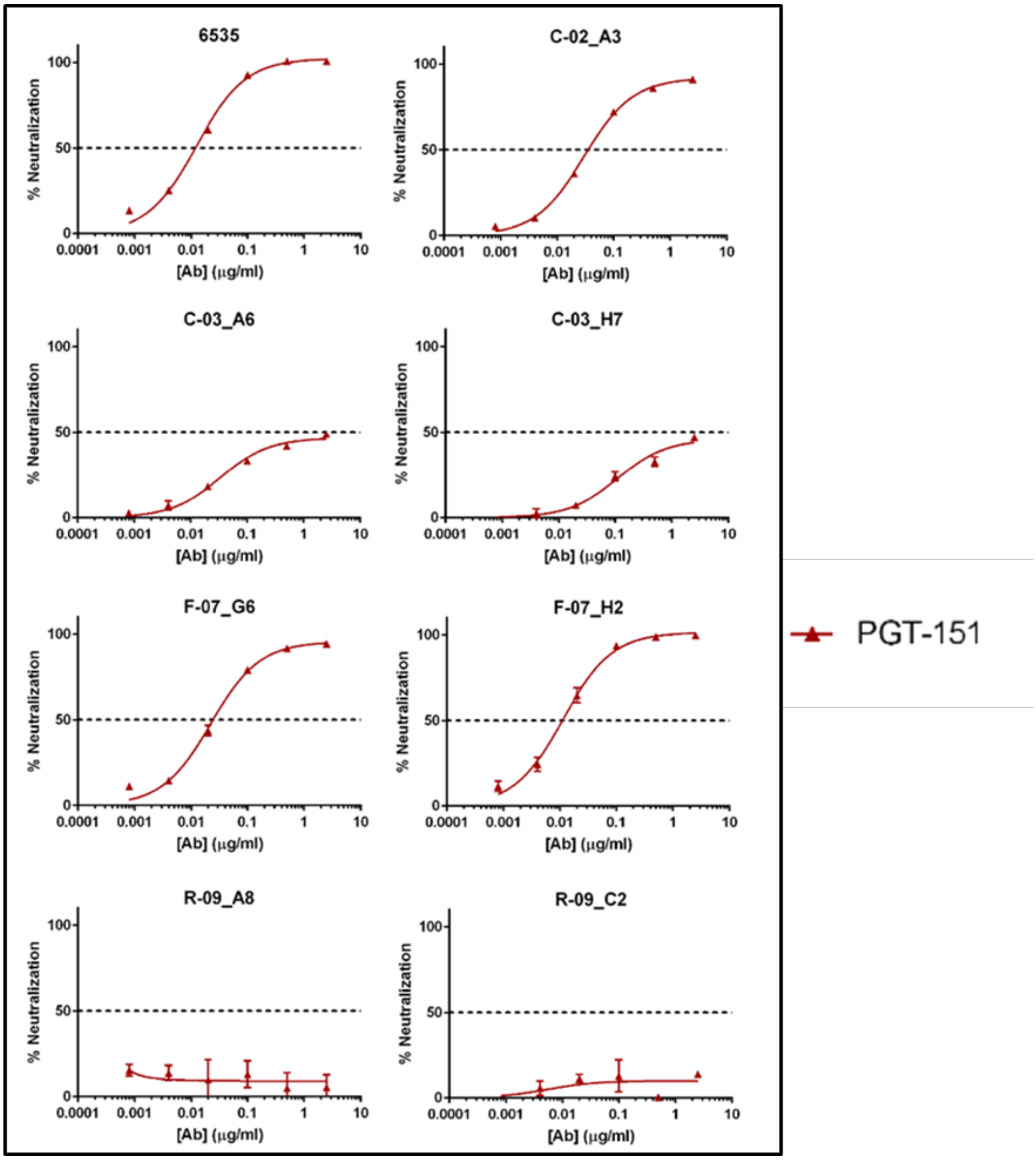

**S6 Fig. Neutralization sensitivities for gp120-gp41 interface monoclonal antibody (mAb) PGT-151.** Donor C-02, C-03, F-07, and R-09 HIV-1 envelopes were expressed as pseudovirions and incubated with gp120-gp41 interface mAb PGT-151 for 1 h then added to TZM-bl target cells. Luciferase was measured after 3 days. Percentage neutralization was calculated relative to cell-only and virus-only controls. The tier 1b subtype B pseudovirus 6535 is used as a neutralization control. Neutralization curves fitted by GraphPad PRISM v10.0.0.

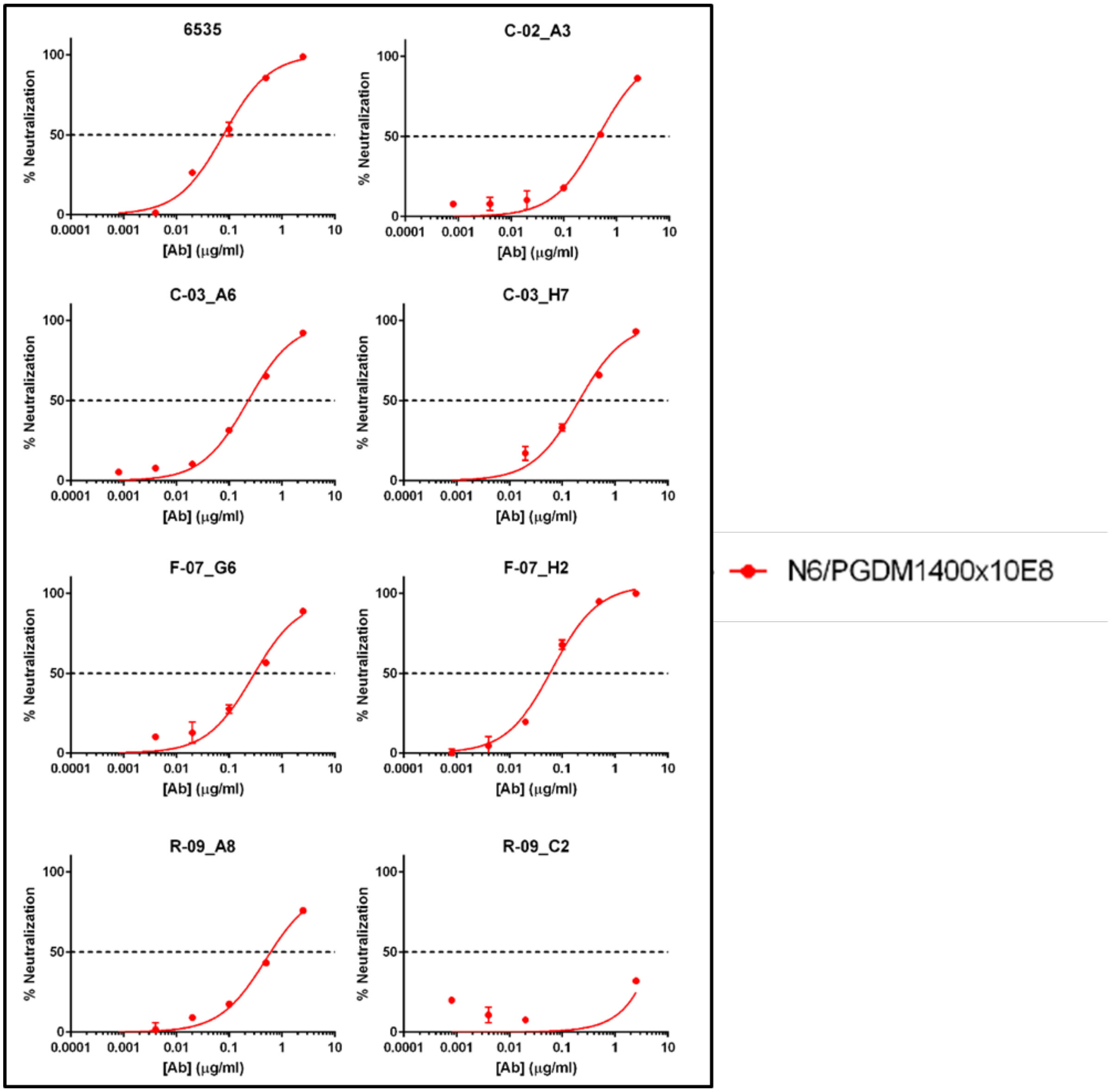

**S7 Fig. Neutralization sensitivities for tri-specific monoclonal antibody (mAb) NG/PGDM1400x10E8.** Donor C-02, C-03, F-07, and R-09 HIV-1 envelopes were expressed as pseudovirions and incubated with tri-specific mAb NG/PGDM1400x10E8 for 1 h then added to TZM-bl target cells. Luciferase was measured after 3 days. Percentage neutralization was calculated relative to cell-only and virus-only controls. The tier 1b subtype B pseudovirus 6535 is used as a neutralization control. Neutralization curves fitted by GraphPad PRISM v10.0.0.

**S1. Quantification of purified autologous immunoglobulin (Ig) isotypes by enzyme-linked immunosorbent assay (ELISA).** Concentrations for purified Ig determined by absorbance at 260nm. Concentrations of purified Ig isotypes (i.e. IgG1-G4, IgA, and IgM) measured by quantitative ELISA. Purified isotype percentages determined by dividing Ig isotype concentration by total Ig concentration and the inputted amounts into neutralization assays calculated.

| **Sample** | **Purified Plasma Ig (mg/mL) [a]** | **IgG (mg/mL) [b]** | **IgG Concentration of Primary Dilution (ug/ml) [c]** | **IgA (mg/mL) [b]** | **IgA Concentration of Primary Dilution (ug/mL) [c]** | **IgM (mg/mL) [b]** | **IgM Concentration of Primary Dilution (ug/mL) [c]** |
|---|---|---|---|---|---|---|---|
| C-02 | 10.824 | 1.942 | 89.7 | 0.154 | 7.1 | 0.50 | 23.1 |
| C-03 | 13.865 | 2.596 | 93.6 | 0.452 | 16.3 | 0.46 | 23.0 |
| F-07 | 8.215 | 1.489 | 90.7 | 0.326 | 19.9 | 0.62 | 37.8 |
| R-09 | 9.747 | 1.939 | 99.5 | 0.370 | 19 | 0.79 | 40.6 |

[a] Purified from plasma total immunoglobulin concentrations (mg/mL) as determined by absorbance at 280nm by Novix DS-11 spectrophotometer.

[b] Purified from plasma, immunoglobulin isotype concentrations (mg/mL) were determined by quantitative human IgG, IgA, and IgM enzyme-linked immunosorbent assay kits (Fisher Scientific, Waltham, MA, USA).

[c] 500ug/mL of total purified immunoglobulins from plasma used to calculate the IgG, IgA, and IgM concentrations of the primary dilution.

**S2 Table. HIV-1 Envelope Mutations in HIV-1 Envs from the non-suppressible viremic donor C-03 that confer resistance to neutralization by b12.**

| | | Potential N-Glycosylation Site Residues [a] | | C3 Domain Residues [b] | | Gp41 Residues [c] | |
|---|---|---|---|---|---|---|---|
| **Donor HIV-1 Envelopes** | **Amino Acid Position197** | **Amino Acid Position 301** | **Amino Acid Position 364** | **Amino Acid Position 369** | **Amino Acid Position 373** | **Amino Acid Position 569** | **Amino Acid Position 675** |
| C-02_A3 | N | N | P | P | M | T | I |
| C-03_A6 | N | N | S | P | M | T | I |
| C-03_H7 | N | N | S | P | M | T | I |
| F-07_G6 | N | N | S | P | M | T | I |
| F-07_H2 | N | N | S | P | M | T | I |
| R-09_A8 | N | N | S | P | T | T | I |
| R-09_C2 | N | N | S | P | T | T | I |

[a] Potential N-Glycosylation Sites (PNGS) N197 and N301 associated with resistance to neutralization by antibody b12. Removal of N197 or N301 increases sensitivity to neutralization.

[b] HIV-1 Env amino acids S364, P369, and T373 are associated with resistance to neutralization by antibody b12. Substitutions S364H, P369L, and T373M associated with increased sensitivity to neutralization by antibody b12.

[c] HIV-1 Env amino acids T569 and I675 are associated with resistance to neutralization by antibody b12. Substitutions T569A and I675V are associated with increased sensitivity to neutralization by antibody b12.

**S3 Table. Discordant Amino Acid Positions between Donor C-03_A6 and C-03_H7 HIV-1 Envelopes.**

| | Nucleotide Position | HIV-1 Env C-03_A6 | HIV-1 Env C-03_H7 | Comment |
|---|---|---|---|---|
| **1** | 318 | E106 | Q106 | Acidic versus amide side chain |
| **2** | 330 | N130 | S130 | Carboxamide versus hydroxyl side chain |
| **3** | 543 | G181 | K181 | Nonpolar versus positively charged side chain |
| **4** | 549 | 183 | N183 | Addition of an extra amino acid |
| **5** | 552 | S184 | N184 | Hydroxyl versus carboxamide side chain |
| **6** | 816 | E272 | D272 | Both have acidic side chains |
| **7** | 885 | V295 | I295 | Both have hydrophobic side chains |
| **8** | 1170 | K390 | Q390 | Positively charged to polar |
| **9** | 1191 | N397 | T397 | Carboxamide versus hydroxyl side chain |
| **10** | 1194 | S398 | N398 | Hydroxyl versus carboxamide side chain |
| **11** | 1197 | *N399 | D399 | Loss of PNGS in C-03_H7 |
| **12** | 1221 | L407 | R407 | Hydrophobic versus positively charged side chain |
| **13** | 1224 | F408 | V408 | Bulky aromatic versus hydrophobic chain |
| **14** | 1230 | *N410 | D410 | Loss of PNGS in C-03_H7 |
| **15** | 1233 | N411 | T411 | Carboxamide versus hydroxyl side chain |
| **16** | 1386 | N462 | K462 | Polar versus positively charged side chain |
| **17** | 1395 | *T465 | *N465 | New PNGS in both that are offset by 2 amino acids |
| **18** | 2430 | Q810 | K810 | Polar versus positively charged side chain |
| **19** | 2463 | L821 | I821 | Both have hydrophobic side chains |
| **20** | 2490 | A830 | T830 | Nonpolar versus polar side chain |
| **21** | 2535 | T845 | I845 | Polar versus hydrophobic side chain |

* Addition or subtraction of potential N-linked glycosylation site.

**S4 Table. HIV-1 Envelope Mutations Associated with PG9, PG16, and PGT-145 Resistance to Neutralization.**

| HIV-1 Envs | Neutralization Profile | Region Spanning Amino Acids 163-176 | Overall Net Charge of Region Spanning Amino Acids 163-176 | Consequence of Amino Acid Position Change |
|---|---|---|---|---|
| CAP45† | Sensitive | TELRDKKQKAYALF | +2 | |
| C-02_A3* | Resistant [a, b] | SGIRDKVKKEYAYF | +2 | |
| C-03_A6 | Resistant [b] | TNIRDKIQKEYALF | +1 | Reduced overall electrostatic charge |
| C-03_H7* | Sensitive | TNIRDKIQKEYALF | +1 | Reduced overall electrostatic charge |
| F-07_G6 | Resistant [a] | SGIRDKVKKEYAYF | +2 | |
| F-07_H2 | Sensitive | SGIRDKVQKEYAYF | +1 | Reduced overall electrostatic charge |
| R-09_A8* | Resistant [a, b, c] | TDMGNKKEKERAFF | +1 | Reduced overall electrostatic charge, but gained PNGS at 128, loss of PNGS at N160, and substitution at Y173R |
| R-09_C2 | Resistant [a, b, c] | TDMNKKEKERAFF | +1 | Reduced overall electrostatic charge, but gained PNGS at 128, loss of PNGS at N160, and substitution at Y173R |

* Repliclone; [a] Resistant to antibody PG9; [b] Resistant to antibody PG16; [c] Resistant to antibody PG145; Residues highlighted in bold green are positively charged; those in bold red are negatively charged.
